# Targeted Alpha Therapy with [^225^Ac]Ac-Macropa-Isatuximab for CD38-positive Hematological Malignancies

**DOI:** 10.1101/2024.11.01.621584

**Authors:** Natalia Herrero Alvarez, David Bauer, Alexa L. Michel, Lukas M. Carter, Jason S. Lewis

**Affiliations:** Department of Radiology and Program in Pharmacology, Memorial Sloan Kettering Cancer Center; New York, NY, USA; Department of Medical Physics, Memorial Sloan Kettering Cancer Center; New York, NY, USA; Departments of Pharmacology and Radiology, Weill Cornell Medicine; New York, NY, USA

**Keywords:** CD38, multiple myeloma, lymphoma, targeted alpha therapy, actinium-225

## Abstract

**Background:** Hematological malignancies include diverse cancers related to immune system cells and blood-forming tissue. Treatment resistance and relapse are common despite the introduction of novel therapeutics. Following the development of the PET probe [^89^Zr]Zr-DFO-isatuximab, targeting the overexpressed CD38 receptor, we developed [^225^Ac]Ac-Macropa-isatuximab for targeted alpha therapy in murine models of human multiple myeloma (MM) and lymphoma.

**Methods:** In vitro studies were performed using the CD38 expressing MM1.S (human MM) and Daudi (human lymphoma). Radiopharmaceutical therapy studies were performed in female NSG mice, and the disseminated disease models were established by intravenous injection of luciferase-transfected cells. Mice were randomized (n = 10 mice per cohort) and received a high activity (5.5 kBq) or low activity (2.7 kBq) of [^225^Ac]Ac-Macropa-Isatuximab in single or multiple cycles. Controls receiving untargeted [^225^Ac]Ac-Macropa-IgG or saline were included. Tumor burden was measured by bioluminescence imaging (BLI). Weight loss greater than 20% or leg paralysis were used as endpoints.

**Results:** [^225^Ac]Ac-Macropa-Isatuximab was obtained with high radiochemical yield and purity (>95%). It displayed high immunoreactivity and excellent stability in human serum over 10 days. The therapeutic effect of [^225^Ac]Ac-mcp-Isatuximab was significant for all the treatment cohorts compared to the saline control after administration of the first cycle. Low activities were better tolerated in general, resulting in extended median survivals. The most successful regimen in the MM model was the dual cycle of low activity, with a median survival of 60 days (P < 0.001). In the lymphoma model, the single high activity of 5.5 kBq and four low activity of 2.7 kBq resulted in similar median survivals of 143 days (P < 0.0001) and 147 days (P < 0.0001). These regimens resulted in complete responses with no detectable cancer cells in some cases. Remarkably, no significant kidney damage was observed in any of the models.

**Conclusion:** [^225^Ac]Ac-Macropa-Isatuximab selectively targets CD38, reducing tumor growth and, in some cases, tumor eradication, especially when applied in multiple cycles. Importantly, it has not been associated with significant toxicities at levels of administered activity required for therapeutic benefit, indicating its promise as a therapeutic option.

## INTRODUCTION

Hematological malignancies comprise a heterogeneous group of lymphoid and myeloid neoplasms that occur due to the dysregulation of normal hematopoietic processes.^1^ They affect the production and function of immune system cells or blood-forming tissue, such as the bone marrow, often leading to fatal outcomes. Clinically, they can be stratified, according to their cell of origin, into three major groups: lymphoma, myeloma, and leukemia. Together, they account for 187,740 cases in the US in 2024, nearly 10% of all new cancer cases diagnosed.^2^

Unfortunately, despite the availability of novel therapeutics, there is a lack of curative approaches, and both treatment resistance and relapse are a common feature for patients with blood cancers. Major challenges when developing therapies for these malignancies include their profound heterogeneity and difficulty achieving and maintaining therapeutic efficacy while sparing progenitor cells.^3^

The use of radiopharmaceuticals for imaging and therapy in oncology has grown extensively over the last few years, especially since the approval of Lutathera and Pluvicto.^4^ In the case of hematological malignancies, targeted radiopharmaceutical therapy represents an advantageous approach to disease management. Most neoplasms are highly sensitive to radiation, and radiopharmaceutical agents hold the potential to overcome resistance and target heterogeneity.

In this context, targeted radioligand therapy, harnessing beta-emitters, has been discussed as a potential treatment.^5, 6^ However, beta therapy is limited by toxicity arising from bone marrow off-targeting^7^. These limitations could be overcome by targeted alpha therapy (TAT), as recently reported.^8-10^ The benefits of alpha particles lie in their shorter tissue path length and higher linear energy transfer (LET), which maximizes their therapeutic efficacy while sparing a higher number of healthy bone marrow precursors.^9, 11^ Specifically, the alpha emitter actinium-225 (^225^Ac), with four net α-particles in its decay chain and 10-day half-life, has emerged as a promising candidate for clinical translation and has recently shown great potential in different preclinical models of hematological malignancies.^12-15^

Moreover, the development of theranostic agents, guided by personalized dosimetry, could allow the pre-selection of patients, suggest the most effective targeted therapies, and measure treatment responsiveness. This would result in interventions based on the individual case.

CD38, a 45 kDa transmembrane glycoprotein, is a well-established cancer hallmark in multiple myeloma (MM) and is currently under investigation in other hematological malignancies. Its expression is correlated with cellular proliferation and disease progression.^16^ Importantly, CD38 is overexpressed in abnormal lymphoid and myeloid cells relative to counterpart healthy cells,^17^ thus an excellent target for theranostics.

We recently reported the preclinical development of [^89^Zr]Zr-DFO-Isatuximab as a highly selective and specific agent for noninvasive CD38-targeted imaging. Its potential was demonstrated in two murine models of CD38+ hematological cancers: an MM and an aggressive type of non-Hodgkin’s lymphoma (NHL).^18^

This work is dedicated to developing a therapeutic counterpart based on the radionuclide ^225^Ac, which was evaluated in our established models. Moreover, we investigated the benefits of a multiple-cycle regime, which could increase treatment efficacy while limiting toxicity.

## MATERIAL AND METHODS

Cell lines, *in vitro* studies, dosimetry, toxicity, hematological analysis, and immunohistochemistry are described in the Supplementary Information.

### TCO Functionalization, ^225^Ac-labeling, and IEDDA Reaction

Antibodies were prepared as an 11 mg/mL solution in PBS and reacted with 20 molar equivalents of NHS-activated trans-cyclooctene (TCO-NHS) at pH 9 for 1 h at 37°C. The functionalized antibody was purified by spin filtration (MW cutoff = 50 kDa, 0.5 mL; Amicon Ultra). Conjugation efficiency was assessed by MALDI-TOF mass spectrometry.

Macropa-PEG_8_-Tetrazine (mcp-PEG_8_-Tz) was synthesized as previously described.^19^ [^225^Ac]Ac(NO_3_)_3_ was purchased from the National Isotope Development Center, reconstituted in hydrochloric acid (0.1 M), and neutralized with 0.25 M NH_4_OAc buffer (0.25 M, pH 5.5) for radiolabeling. After the addition of the precursor (10 nmol, in 10 μL DMSO), the reaction was incubated for 5 min at 37°C, and the radiochemical conversion (RC) was determined by instant thin-layer chromatography (iTLC) at secular equilibrium using 50 mM ethylenediaminetetraacetic acid (EDTA) as the mobile phase. The resulting [^225^Ac]-mcp-PEG_8_-Tz was reacted with the corresponding mAb-TCO for 5 min at 37°C. Completion of the inverse-electron demand Diels−Alder reaction was confirmed by iTLC using a mobile phase of NH_4_OAc (pH 5.5, 0.25 M) and methanol (40:60). In this system unconjugated [^225^Ac]Ac-mcp-PEG_8_-Tz moves along with the solvent front.

### Animal Models and Bioluminescence Imaging

All animal experiments performed in this study were approved by the Institutional Animal Care and Use Committee and Research Animal Resource Center at Memorial Sloan Kettering Cancer Center. *In vivo* studies were performed in female NSG mice (NOD.Cg-Prkdcscid Il2rgtm1Wjl/SzJ; 6–8 weeks old; Jackson Laboratories, Bar Harbor, Maine). The disseminated disease models were established by intravenous injection (IV) of MM.1S-Luc (1 × 10^6^ cells) and Daudi-Luc cells (0.5 × 10^6^ cells). Tumor burden was monitored by bioluminescence imaging (BLI) with an IVIS Spectrum-CT (PerkinElmer, Melville, NY). Whole-body regions of interest were used for quantification, and tumor burden was expressed as total flux (p/s).

### Radiopharmaceutical Therapy Studies

Radiopharmaceutical therapy studies were designed to evaluate dose dependence by administering a high activity of 5.5 kBq and a low activity of 2.7 kBq. Additionally, we investigated the impact of a single administration versus administration in multiple cycles. Administered activities were guided by dosimetry estimates and literature data.^9, 14, 20^ Mice were randomized based on bioluminescence imaging (BLI) 9–12 days after inoculation into 6 arms (n = 10), including two control cohorts, one receiving saline and the second receiving 5.5 kBq of [^225^Ac]Ac-mcp-IgG, a control immunoconjugate lacking specific targeting. Two arms were treated with single cycles of [^225^Ac]Ac-mcp-Isatuximab, 5.5 kBq or 2.7 kBq, while the other two received a second cycle of the same activity for the MM model, and four cycles for the lymphoma model. The arms receiving [^225^Ac]Ac-mcp-Isatuximab were coinjected with 500 μg of isotype IgG for Fc blocking. The specific activity of the formulation was adjusted such that 30 μg of antibody was administered for each cycle. Mice were monitored by BLI weekly, and toxicity was evaluated by monitoring overall condition, body weight loss, and retroorbital blood draws biweekly. Endpoints were defined as weight loss greater than 20% or leg paralysis; otherwise, the study was terminated at day 150. After reaching the endpoint, selected mice were fixed in formalin and submitted to a Laboratory of Comparative Pathology for evaluation.

### Statistical Analysis

Statistical analyses were performed using GraphPad Prism software version 10. Statistical significance was evaluated with a t-test, a one-way ANOVA test, or a two-way ANOVA test, as appropriate for the number of variables, and the threshold for statistical significance was set at *p* < 0.05. Data presented are expressed as mean ± SD.

### Data Availability

All data generated during this study are included in this published article and its supplementary information files, or otherwise available upon request from the corresponding author.

## RESULTS

### [^225^Ac]Ac-mcp-isatuximab Synthesis, Immunoreactivity and Stability

[^225^Ac]Ac-mcp-isatuximab was prepared in 2-steps (Fig S1) using an inverse-electron demand Diels−Alder reaction to click isatuximab-TCO with [^225^Ac]Ac-mcp-PEG_8_-Tz. Radiolabeling of mcp-PEG_8_-Tz proceeded with high radiochemical conversion (> 99%), and upon incorporation onto the antibody, [^225^Ac]Ac-mcp-isatuximab was obtained with > 95% purity (Fig S2). An average of 7 TCO moieties were attached per antibody molecule during the bioconjugation (Fig S3). The resulting radioimmunoconjugate displayed a high immunoreactivity and maintained high specific binding to the MM.1S (derived from human multiple myeloma) and Daudi (derived from human lymphoma) cell lines, as well as excellent stability in human serum over 10 days (Fig S4).

### Animal Models and Dosimetry

Evaluation of [^225^Ac]Ac-mcp-isatuximab was conducted in disseminated murine models of MM and NHL, benchmarks of CD38+ hematological malignancies. These models, established in NGS mice upon intravenous injection of the cell lines, allow tumor cells to circulate and infiltrate in the BM and other hematopoietic organs, being more consistent with the real pathology of these malignancies. It is important to note that these models present extensive extramedullary disease and, therefore, resemble very aggressive or late-stage disease, particularly the lymphoma model.^21^ Adopting the theranostic approach commonly considered in personalized nuclear medicine, we utilized the biodistribution data obtained with the imaging counterpart, [^89^Zr]Zr-DFO-Isatuximab, to project dosimetry estimates and select administered activity levels for radiopharmaceutical therapy with [^225^Ac]Ac-Macropa-isatuximab. The dosimetry estimates implicated the hematopoietic (red) marrow and liver as potential dose-limiting organs for therapy with [^225^Ac]Ac-Macropa-isatuximab. The RBE-weighted dose coefficient for uninvolved red marrow was estimated from the activity concentration in blood^22^ to be 0.14 Gy-eq/kBq in the MM.1S model and 0.56 Gy-eq/kBq in the Daudi model. The liver received 3.2 Gy-eq/kBq (MM1.S model) and 1.4 Gy-eq/kBq (Daudi model). RBE-weighted doses for the tumor tissue/involved marrow were 8.7 Gy-eq/kBq in the MM.1S model and 4.2 Gy-eq/kBq in the Daudi model, estimated from the marrow samples harvested from the distal femur or obtained from contours of diseased sites in the lumbar vertebrae obtained from PET images. Therapeutic indices (i.e., tumor-to-normal tissue dose ratios) for the uninvolved marrow were 63 and 8 in the MM.1S and Daudi models, respectively, while those for the liver were 2.7 and 3.0. See Supplemental Table S5B for tabulated dose coefficients for additional tissues.

### Radiopharmaceutical Therapy with [^225^Ac]Ac-mcp-Isatuximab in MM model

The therapy study in mice xenografted with the MM.1S tumor model started 9 days after cell injection when mice were randomized into the different treatment and control cohorts (schematized in Fig 1A). The therapeutic effect of [^225^Ac]Ac-mcp-Isatuximab was significant for all the treatment cohorts compared to the saline control after administration of the first cycle, as determined by the tumor burden measurements obtained from the BLI at day 28 (Fig 1B). Tumor reduction was dose-dependent, with the high administered activity of 5.5 kBq producing a superior delay of tumor growth than the 2.7 kBq administration. Administration of the second cycle on day 29 also had a significant therapeutic effect and further delayed the tumor progression, as demonstrated by BLI measurements on day 43 (Fig 1C).

**Figure 1.**
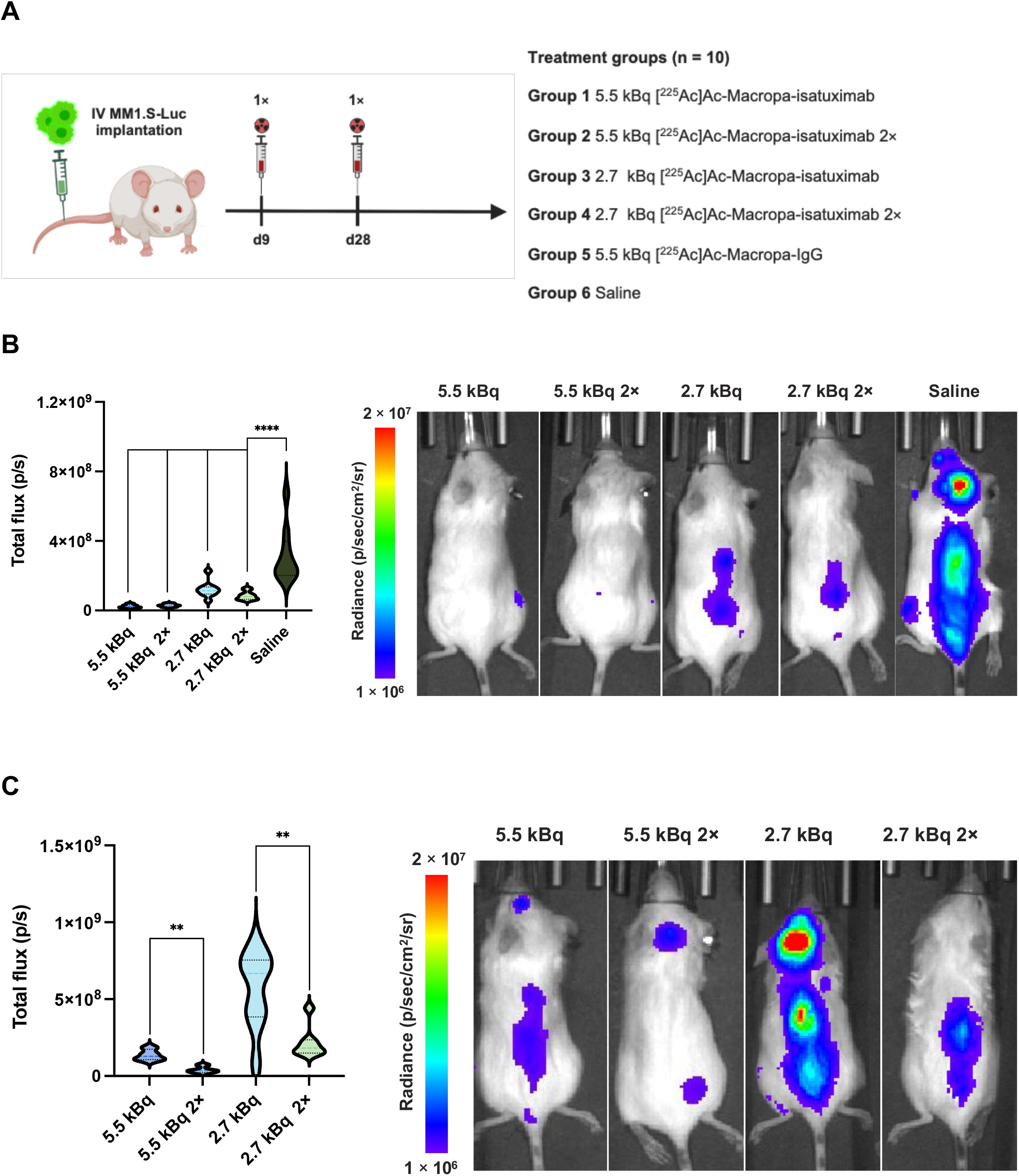
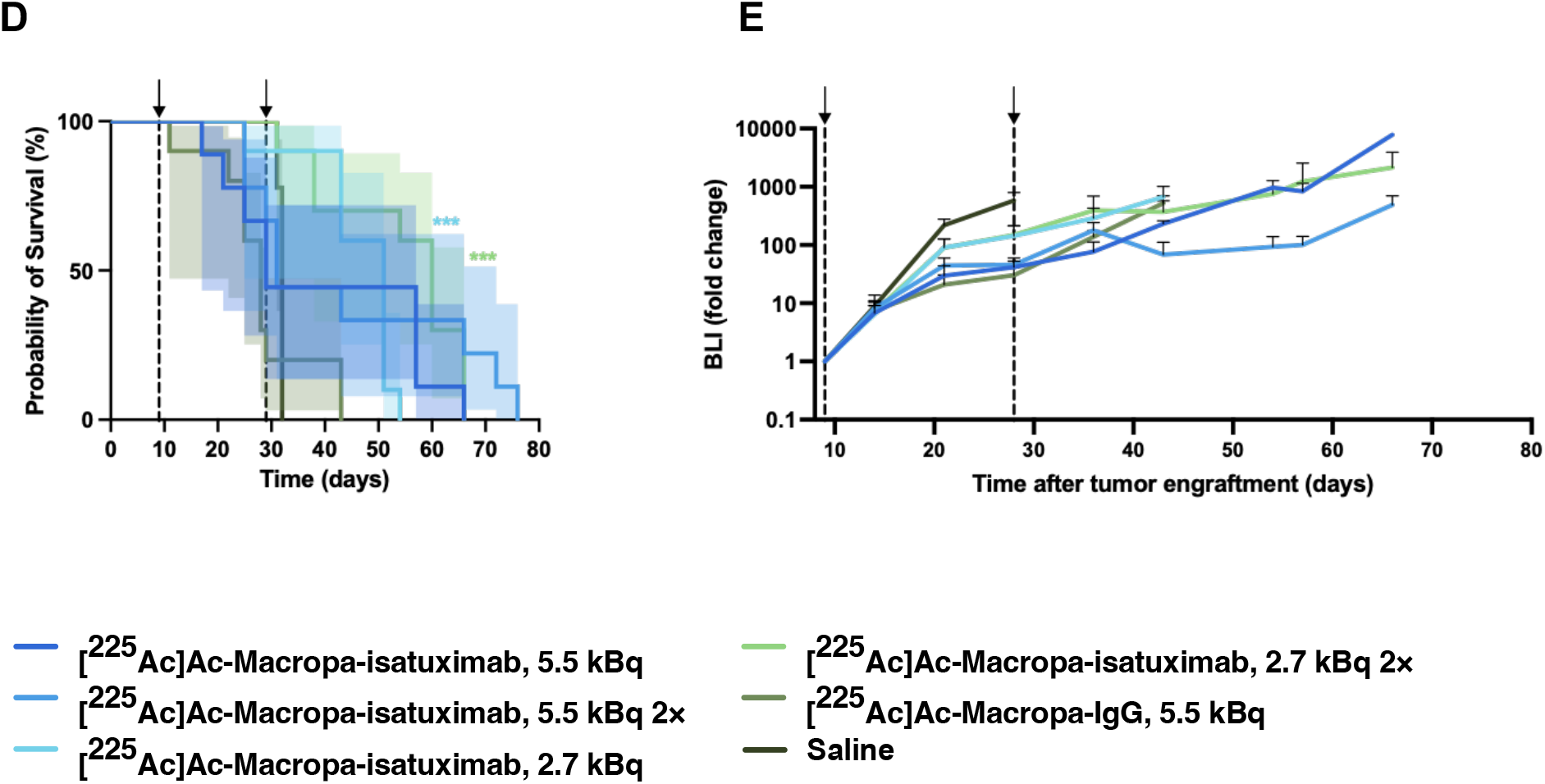
Radiopharmaceutical Therapy Study in Multiple Myeloma Model. Schematic of the therapy study in the MM.1S model (A). Activity-dependence (B) and the tumor response (C) are represented by quantification of the tumor burden measured by BLI. All the treatment groups showed significant tumor growth inhibition compared to the saline group at day 28. Administration of two cycles yielded significantly more inhibition than a single cycle, both at 5.5 kBq and 2.7 kBq, as seen on day 43. Probability of survival (D) and plot of fold change of tumor burden as measured by BLI (E) of NSG mice with MM.1S disseminated disease. Significant probability of survival was observed in 2.7 kBq [^225^Ac]Ac-mcp-isatuximab, and 2.7 kBq [^225^Ac]Ac-mcp-isatuximab 2× compared to saline and IgG isotype control, yielding a median survival of 51 and 60 days, respectively. Shading represents 95% Cl. Gridlines indicate the day(s) administration. **P < 0.01, ***P < 0.001, ****P < 0.0001.

However, toxicity was also dose-dependent, and while the high administered activities, administered via single- or dual-cycle schedule, produced a greater tumor control, they resulted in decreased median survival as determined by the Kaplan-Meier curve (Fig 1D) of 29 days and 31 days, respectively, with no statistical difference between them. Weight loss was the main endpoint reached in these cohorts (Fig S9). In contrast, the low activity was well tolerated, and the administration of additional cycles further sustained the delay of tumor growth, resulting in the most successful treatment, with a median survival of 51 days (P < 0.001) for single and 60 days (P < 0.001) for dual cycle treatments. No limiting toxicities were observed was associated with the low activity administrations and the endpoint of these cohorts was disease progression. Reduction of tumor in the cohort receiving untargeted [^225^Ac]Ac-mcp-IgG was associated with total bone marrow ablation, which resulted in extensive whole-body toxicity and the lowest median survival (28 days). Untreated control mice were euthanized on day 32 due to limb paralysis. The treatment study was finalized on day 72. Tumor progression over time normalized to baseline disease (Fig 1E) shows transient plateaus after each dose administration compared to subsequent exponential growth. This suggests that a higher frequency of low activity cycles could increase the overall survival.

Additionally, the acquired blood panels (Fig S9) supported these observations, showing dose-dependent and transitory hematologic toxicity for treatment cohorts, and severe toxicity for the untargeted cohort, particularly toward white blood cells and platelets.

Finally, selected mice were submitted for a comprehensive pathological evaluation. Hematoxylin- and eosin-stained (H&E) vertebral sections (Fig 2), exposing BM composition, evidenced the efficacy and specificity of [^225^Ac]Ac-mcp-isatuximab, in accordance with the previously discussed results. A single high activity cycle (Fig 2A) greatly limited the growth of neoplastic cells. In contrast to saline (Fig 2B) and untargeted controls, it maintained a healthy BM configuration, the latter showing the mentioned BM ablation (Fig 2C). A higher presence of neoplastic cells is observed for the low activity single-cycle treatment (Fig S11, S12). Only the ovaries presented treatment-associated toxicity, which could be partially explained by their high radiosensitivity and relatively close local proximity. Remarkably, no significant kidney damage was observed (Fig S12). These results indicate that multiple administration of low activity is a promising approach to surpassing toxicity, and that administering several cycles at a high frequency (1–2 week intervals) may further prolong the overall survival.

**Figure 2.**
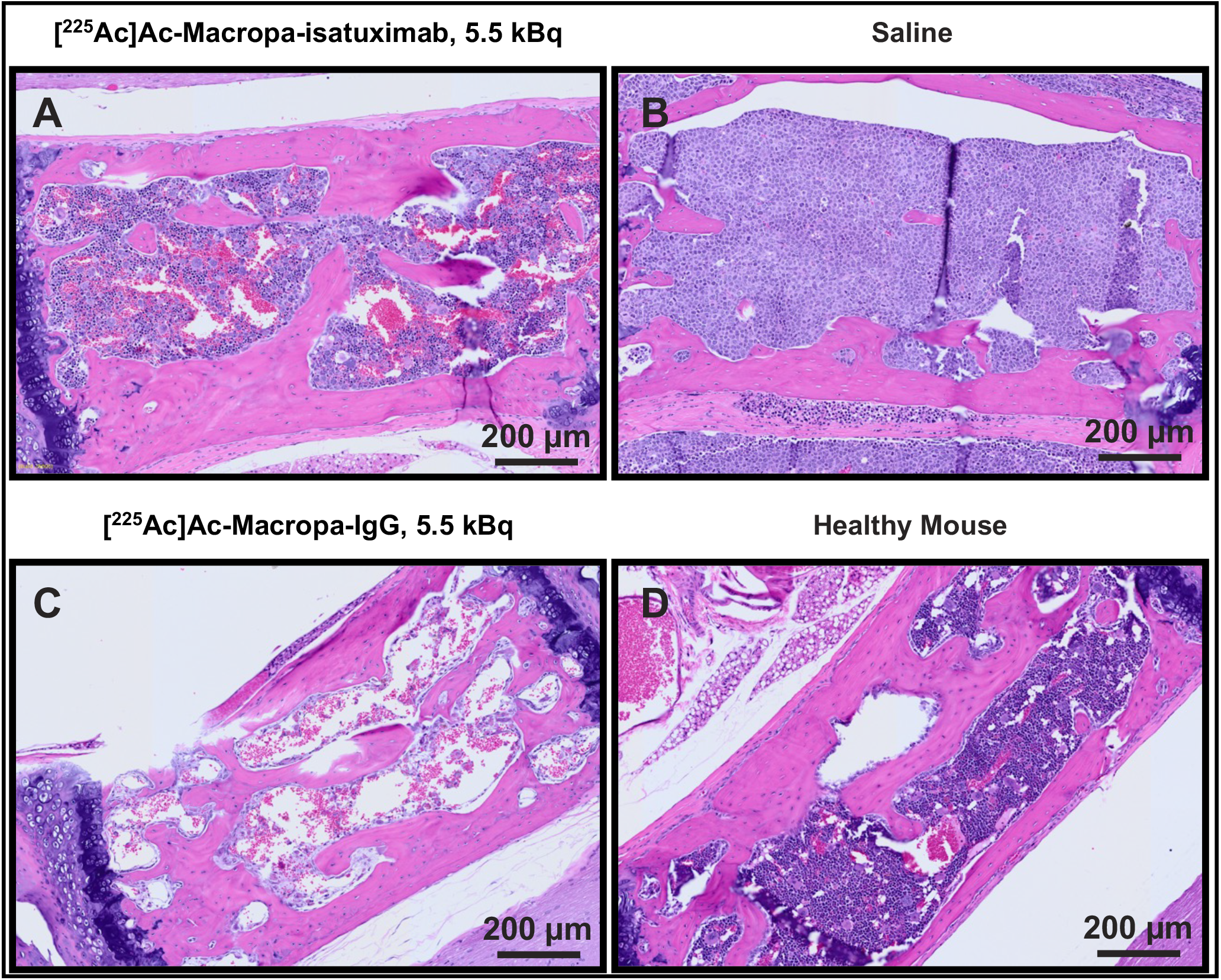
Histology of Hematoxylin- and Eosin-Stains of the MM.1S Model. Representative stained vertebral sections of MM.1S disseminated NSG mice (A–C) and one healthy mouse as a reference (D). Mice treated with 5.5 kBq [^225^Ac]Ac-mcp-isatuximab (A), saline control (B), untargeted 5.5 kBq [^225^Ac]Ac-mcp-IgG (C). Histopathology revealed minimal presence of neoplastic cells and a high number of healthy erythroid and myeloid precursors in treated mice (A). Saline mice showed no presence of healthy precursors, the tumor cells have replaced the bone marrow and infiltrated into the surrounding tissues (B). On the other hand, mice administered with untargeted 5.5 kBq [^225^Ac]Ac-mcp-IgG presented severe hypoplasia and hemorrhage due to bone marrow destruction (C). Bone marrow comprises healthy erythroid and myeloid precursors in healthy mice (D).

### Radiopharmaceutical Therapy with [^225^Ac]Ac-Macropa-Isatuximab in NHL model

The minimal toxicity and moderate tumor response observed for the MM model indicated that increasing the number of cycles could be a promising approach. This would be particularly relevant for the higher aggressiveness of the NHL model. The NHL treatment study was designed with four cycles with weekly administrations, starting at day 12 post-tumor inoculation (schematized in Fig 3A). All treatment arms demonstrated a rapid decrease in tumor burden after the first administration in contrast to untreated mice, which had to be euthanized at day 36 due to high limb paralysis (Fig 3B). Mice receiving multiple high activity cycles presented pronounced toxicity manifested as weight loss, which was the endpoint for 9 out of 10 mice and resulted in a median survival of 33 days. The surviving mouse was euthanized by the end of the study, on day 150, without apparent signs of disease. The single low activity cycle (1 × 2.7 kBq) did not display evident toxicity and did not show disease as measured by BLI for several weeks. Upon disease recurrence, the tumor burden increased rapidly, and mice had to be euthanized due to disease progression, presenting a median survival of 89 days (P < 0.0001). The pronounced difference in individual tumor responses (Fig S14) when using the low activity strongly indicates the importance of personalized dosimetry. Cohorts receiving a single high activity of 5.5 kBq and 4 low activity of 2.7 kBq (group 1 and group 4) resulted in similar median survivals of 143 days (P < 0.0001) and 147 days (P < 0.0001), respectively. Most importantly, mice from these cohorts experienced a complete response by BLI, with 50% surviving until the end of the study (Fig 4).

**Figure 3.**
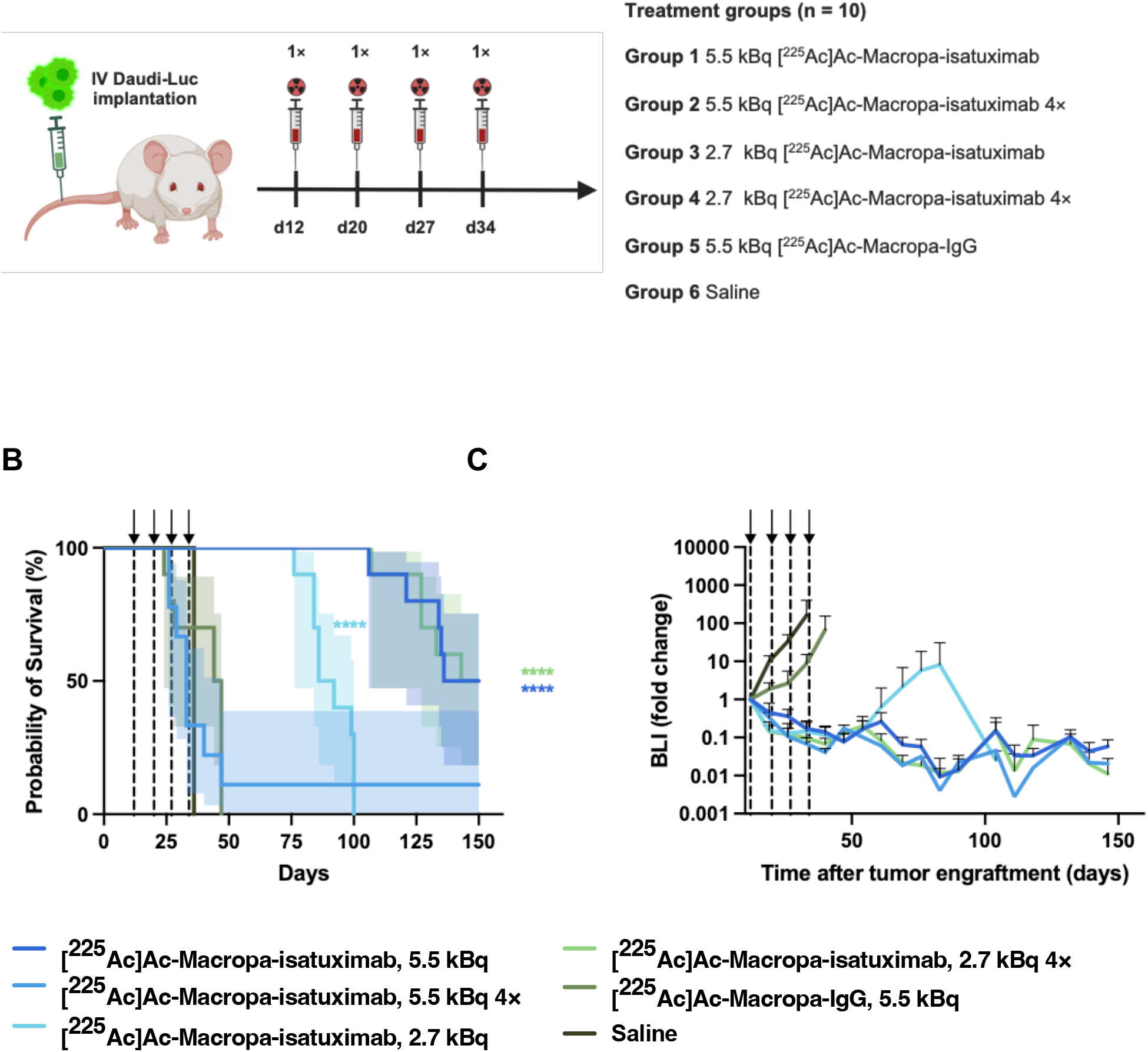
Radiopharmaceutical Therapy Study in Lymphoma Model. Schematic of the therapy study in the Daudi model (A). Probability of survival (B) and fold change of tumor burden as measured by BLI (C) of NSG mice with Daudi disseminated disease. Significant probability of survival was observed in 5.5 kBq [^225^Ac]Ac-mcp-isatuximab, 2.7 kBq [^225^Ac]Ac-mcp-isatuximab, and 2.7 kBq [^225^Ac]Ac-mcp-isatuximab 4× compared to saline and IgG isotype control, yielding a median survival of 168, 89 and 171.5 days, respectively. A high therapeutic response is demonstrated by 5.5 kBq [^225^Ac]Ac-mcp-isatuximab and 2.7 kBq [^225^Ac]Ac-mcp-isatuximab 4×, irradicating the tumor burden. Tumor relapse was not observed in the 5.5 kBq [^225^Ac]Ac-mcp-isatuximab cohort. Therefore, this dosing regime can be considered curative. Shading represents 95% Cl. Gridlines indicate the dose administration. ****P < 0.0001.

**Figure 4.**
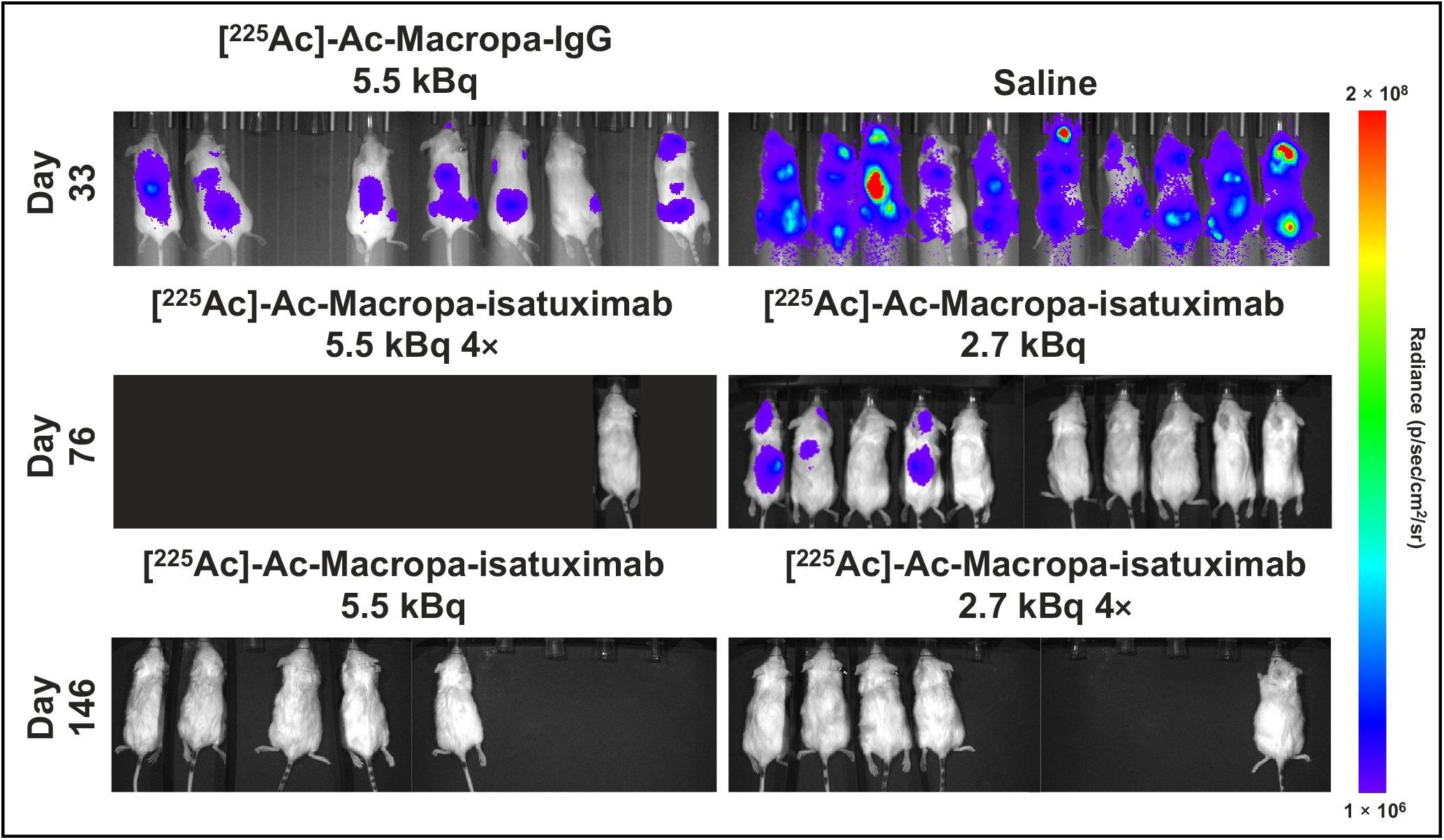
BLI of NSG mice with Daudi disseminated disease. All cohorts of the therapy study are represented. Images shown were obtained near the termination day of each specific cohort. Cohorts receiving 5.5 kBq [^225^Ac]Ac-mcp-isatuximab and 2.7 kBq [^225^Ac]Ac-mcp-isatuximab 4x experienced a complete response.

Additionally, complete blood cell count revealed only mild hematologic toxicity for single 5.5 kBq administration and 2.7 kBq, both single and multiple cycle administration. Radiotoxicity was pronounced for multiple 5.5 kBq and untargeted doses, but still reversible (Fig S16).

Lastly, the complete pathology study confirmed the therapeutic efficacy of [^225^Ac]Ac-mcp-isatuximab in this disseminated model of NHL, with optimal administration regimens of single-cycle 5.5 kBq and multiple-cycle 2.7 kBq. Hematoxylin- and eosin-stained (H&E) femur sections (Fig 5) showed no presence of cancer cells in these cohorts (Fig 5A and Fig 5C) and, more importantly, no long-term bone marrow damage associated with the treatment. Disease recurrence is observed in the cohort receiving the single low dose (Fig 5B). Untreated mice (Fig 5D) and those who relapsed presented tumor infiltration in multiple organs, recapitulating features of clinically aggressive NHL. We had previously observed these features during imaging studies. No adverse renal effects were detected, and only ovarian atrophy was considered a direct result of the treatment (Fig S18).

**Figure 5.**
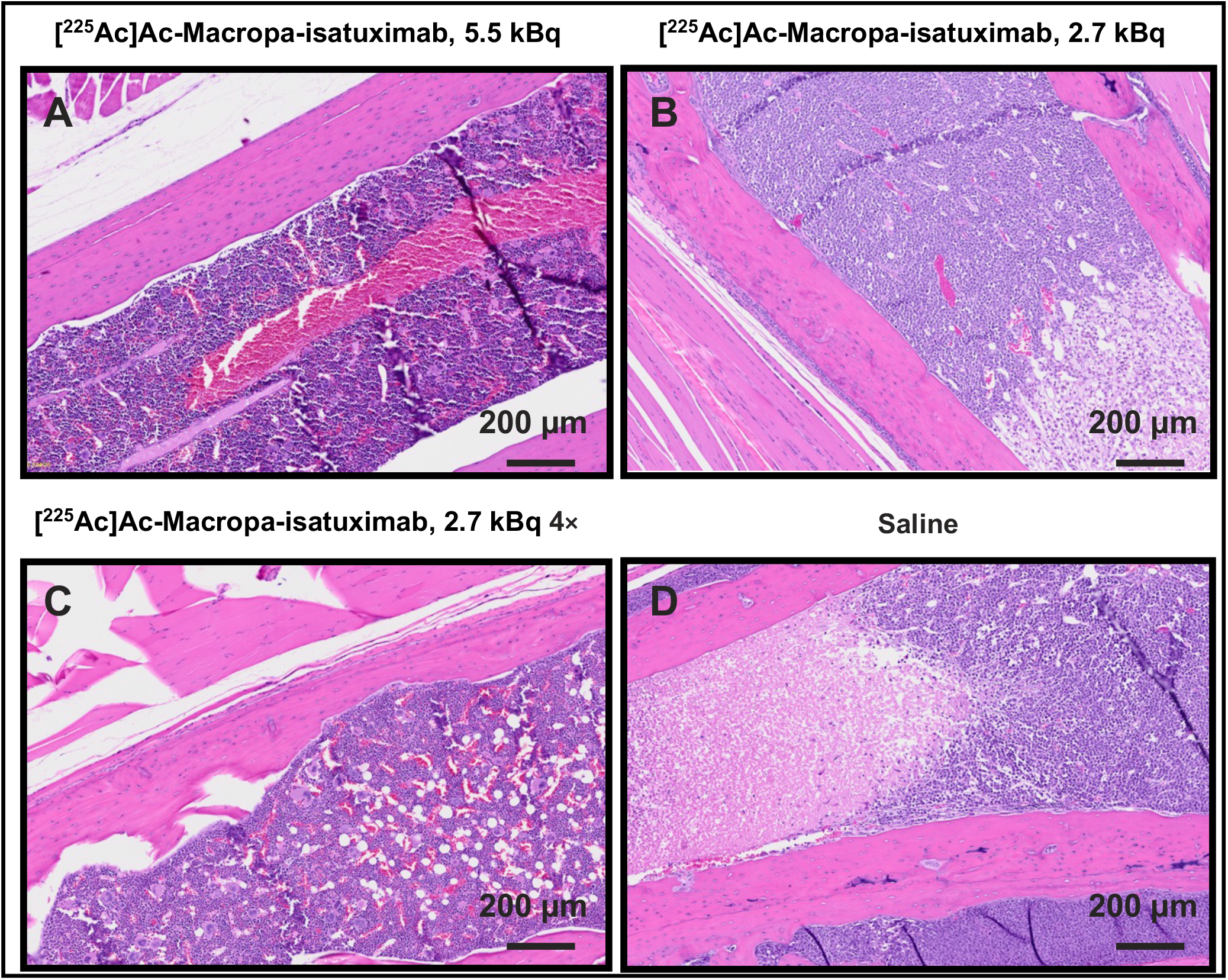
Histology of Hematoxylin- and Eosin-Stains of the Daudi Model. Representative femur sections of Daudi disseminated NSG mice (A–D). Mice treated with [^225^Ac]Ac-mcp-isatuximab: 5.5 kBq (A), 2.7 kBq (B), 2.7 kBq 4× (C) and saline control (D). Histopathology revealed no neoplastic cells in mice treated with 5.5 kBq and 2.7 kBq 4× regimens (A and C); therefore, these mice are considered complete responders. Mice treated with a 2.7 kBq (B) single cycle showed neoplastic round cell infiltration with occasional tumor necrosis due to disease recurrence. Saline control mice (D) presented neoplastic round cell infiltration with frequent tumor necrosis consistent with a high degree of disease.

The holistic pathological evaluation of the treated mice also revealed an unexpected finding. Two mice, from single 5.5 kBq and the 4 × 2.7 kBq regimens, showed tumor burden, which was not detected via BIL since the cells were no longer expressing the luciferase reporter gene (Fig S19). However, flow cytometry of the Daudi cell lines demonstrated stable gene transduction (>95%). This could indicate that some cancer cells have lost the reporter gene *in vivo* or that a very small subclass of not-transduced cells had unmatched growth characteristics and targeting properties. This is an explicit limitation of murine disseminated models and highlights the importance of better diagnostic tools during treatment to determine therapy effectiveness. However, these limitations in BLI detection did not interfere with the therapy survival studies since endpoints included the well-being, weight loss and changes in the blood panel of all mice.

## DISCUSSION

In this work, we presented the synthesis, characterization, and systematic evaluation of [^225^Ac]Ac-mcp-isatuximab in disseminated murine models of MM and NHL as a therapeutic agent for CD38-expressing malignancies.

We used our optimized 2-step click-based approach to prepare the radioimmunoconjugate with high radiochemical yield and purity. [^225^Ac]Ac-mcp-isatuximab displayed high immunoreactivity and serum stability.

MM remains incurable despite the introduction of new treatments, largely due to the development of resistance and the lack of optimal imaging methods. The *in vivo* results demonstrated the capacity of [^225^Ac]Ac-mcp-isatuximab to selectively kill CD38+ tumor cells while sparing healthy bone marrow precursors due to the short path length of alpha particles.

We evaluated the efficacy of this radiopharmaceutical therapy in the challenging MM and lymphoma models on the grounds of clinical relevance. The MM.1S and Daudi models mimic the disseminated nature of multiple myeloma and NHL, respectively. Unlike traditional xenograft models, which fail to replicate the widespread systemic involvement seen in human disease, these models present a more realistic scenario for studying the behavior of these cancers and their responses to systemic radiopharmaceutical therapy. This is especially important in evaluating treatments that rely on targeting multiple tumor sites simultaneously, providing insights that are more translatable to clinical settings. However, despite these strengths, these models also present limitations. The use of highly immunodeficient NSG mice, necessary for the propagation of human MM.1S/Daudi cells, introduces an unrealistic level of radiosensitivity vis a vis radiogenic healthy tissue toxicity; NSG mice lack critical DNA repair mechanisms and immune system components, making them far more susceptible to radiogenic toxicity than human patients. As a result, the maximum tolerated activity and maximal therapeutic benefit in these studies may be underestimated.

Despite this limitation, [^225^Ac]Ac-mcp-isatuximab significantly curtailed myeloma disease progression in a dose-dependent manner, and administration of multiple cycles of low activity sustained the therapeutic activity. This suggests that it may be optimal to begin with a high initial activity cycle followed by multiple low activity cycles with a personalized frequency to optimize effectiveness. Moreover, for most of the tested administration regimens, anticancer efficacy was achieved without substantial toxicity to normal organ systems, and normal organ RBE-weighted doses remained within accepted limits for normal organ absorbed dose established for external beam radiotherapy (Fig S5).^23^

Subsequently, we evaluated [^225^Ac]Ac-mcp-isatuximab in an NHL model, introducing a higher number and frequency of therapy cycles. Although more aggressive, this disease model is also more radiosensitive; thus, a higher response was observed. A pronounced reduction in tumor burden was observed after the first cycle, and the remission was sustained over time in a dose-dependent manner. In fact, the treatment was curative in most animals that received one activity cycle or multiple low activity cycles. Animals that received a single low dose relapsed at some point in the study and showed pronounced differences in individual responses. This advocates for personalized image-guided treatment and disease monitoring in this malignancy.

Overall, this study supports the advantageous features of targeted alpha therapy for personalized treatment of hematological malignancies, particularly if used as a theranostics pair for disease management. Moreover, no severe toxicity was associated with this therapy. Certainly, results suggest clinical translation should be prioritized as the next step, given that the limitations of murine models likely preclude further meaningful optimization or the potential to obtain additional valuable insights. Based on our findings, targeted alpha therapy against CD38 could be a life-saving strategy for patients who have exhausted all alternative treatment options.

## CONCLUSIONS

[^225^Ac]Ac-mcp-isatuximab displays excellent physicochemical and *in vitro* characteristics. It selectively targets CD38, reducing tumor growth and, in some cases, tumor eradication, especially when applied in multiple cycles. Importantly, it has not been associated with significant toxicities at levels of administered activity required for therapeutic benefit, indicating its promise as a therapeutic option.

## ACKNOWLEDGMENTS

We would like to acknowledge the support of the Radiochemistry and Molecular Imaging Probes Core Facility, the Small Animal Imaging Facility, the Molecular Cytology Core Facility, and the Tri-Institutional Laboratory of Comparative Pathology.

## KEY POINTS

### QUESTION

Can [^225^Ac]Ac-Macropa-isatuximab be an effective therapeutic counterpart of previously developed [^89^Zr]Zr-DFO-isatuximab for targeted alpha therapy in disseminated models of CD38+ hematological malignancies?

### PERTINENT FINDINGS

[^225^Ac]Ac-Macropa-isatuximab could be prepared with high radiochemical yield and purity, maintaining high stability. The radioimmunoconjugate displayed great specificity *in vitro* and *in vivo*. CD38-targeted alpha therapy successfully slowed tumor growth in a murine model of disseminated human MM and showed curative efficacy in a murine model of disseminated human NHL.

### IMPLICATIONS FOR PATIENT CARE

[^225^Ac]Ac-Macropa-isatuximab holds excellent potential for targeted radioimmunotherapy of patients preselected by CD38-targeted PET imaging. The lack of treatment-acquired resistance associated with RIT and the ability to determine personalized regimes with the imaging counterpart hold great clinical relevance.

## REFERENCES

1. Hu, D.; Shilatifard, A., Epigenetics of hematopoiesis and hematological malignancies. Genes Dev 2016, 30 (18), 2021–2041.

2. Siegel, R. L.; Giaquinto, A. N.; Jemal, A., Cancer sta7s7cs, 2024. CA: A Cancer Journal for Clinicians 2024, 74 (1), 12–49.

3. Jiang, Y.; Lin, W.; Zhu, L., Targeted Drug Delivery for the Treatment of Blood Cancers. Molecules 2022, 27 (4), 1310.

4. Lapi, S. E.; Scott, P. J. H.; Scott, A. M.; Windhorst, A. D.; Zeglis, B. M.; Abdel-Wahab, M.; Baum, R. P.; Buatti, J. M.; Giammarile, F.; Kiess, A. P.; Jalilian, A.; Knoll, P.; Korde, A.; Kunikowska, J.; Lee, S. T.; Paez, D.; Urbain, J.-L.; Zhang, J.; Lewis, J. S., Recent advances and impending challenges for the radiopharmaceutical sciences in oncology. The Lancet Oncology 2024, 25 (6), e236–e249.

5. Barbet, J.; Bardiès, M.; Bourgeois, M.; Chatal, J.-F.; Chérel, M.; Davodeau, F.; Faivre-Chauvet, A.; Gestin, J.-F.; Kraeber-Bodéré, F., Radiolabeled Antibodies for Cancer Imaging and Therapy. In AnBbody Engineering: Methods and Protocols, Second EdiBon, Chames, P., Ed. Humana Press: Totowa, NJ, 2012; pp 681–697.

6. Duray, E.; Lejeune, M.; Baron, F.; Beguin, Y.; Devoogdt, N.; Krasniqi, A.; Lauwers, Y.; Zhao, Y. J.; D’Huyvetter, M.; Dumoulin, M.; Caers, J., A non-internalised CD38-binding radiolabelled single-domain antibody fragment to monitor and treat multiple myeloma. Journal of Hematology & Oncology 2021, 14 (1), 183.

7. Visser, O. J.; Perk, L. R.; Zijlstra, J. M.; van Dongen, G. A.; Huijgens, P. C.; van de Loosdrecht, A. A., Radioimmunotherapy for indolent B-cell non-Hodgkin lymphoma in relapsed, refractory and transformed disease. BioDrugs 2006, 20 (4), 201–7.

8. Fichou, N.; Gouard, S.; Maurel, C.; Barbet, J.; Ferrer, L.; Morgenstern, A.; Bruchertseifer, F.; Faivre-Chauvet, A.; Bigot-Corbel, E.; Davodeau, F.; Gaschet, J.; Chérel, M., -Dose An7-CD138 Radioimmunotherapy: Bismuth-213 is More Efficient than Lutetium-177 for Treatment of Multiple Myeloma in a Preclinical Model. FronBers in Medicine 2015, 2.

9. Minnix, M.; Adhikarla, V.; Caserta, E.; Poku, E.; Rockne, R.; Shively, J. E.; Pichiorri, F., Comparison of CD38-Targeted α-Versus β-Radionuclide Therapy of Disseminated Multiple Myeloma in an Animal Model. Journal of Nuclear Medicine 2021, 62 (6), 795–801.

10. Cicone, F.; Santo, G.; Bodet-Milin, C.; Cascini, G. L.; Kraeber-Bodéré, F.; Stokke, C.; Kolstad, A., Radioimmunotherapy of Non-Hodgkin B-cell Lymphoma: An update. Seminars in Nuclear Medicine 2023, 53 (3), 413–425.

11. Tranel, J.; Feng, F. Y.; James, S. S.; Hope, T. A., Effect of microdistribution of alpha and beta-emitters in targeted radionuclide therapies on delivered absorbed dose in a GATE model of bone marrow. Phys Med Biol 2021, 66 (3), 035016.

12. Dawicki, W.; Allen, K. J. H.; Jiao, R.; Malo, M. E.; Helal, M.; Berger, M. S.; Ludwig, D. L.; Dadachova, E., Daratumumab-(225)Actinium conjugate demonstrates greatly enhanced an7tumor ac7vity against experimental mul7ple myeloma tumors. Oncoimmunology 2019, 8 (8), 1607673.

13. Kinder, M.; Bahlis, N. J.; Malavasi, F.; De Goeij, B.; Babich, A.; Sendecki, J.; Rusbuldt, J.; Bellew, K.; Kane, C.; Van de Donk, N., Comparison of CD38 antibodies in vitro and ex vivo mechanisms of ac7on in mul7ple myeloma. Haematologica 2021, 106 (7), 2004–2008.

14. Wadhwa, A.; Wang, S.; Patiño-Escobar, B.; Bidkar, A. P.; Bobba, K. N.; Chan, E.; Meher, N.; Bidlingmaier, S.; Su, Y.; Dhrona, S.; Geng, H.; Sarin, V.; VanBrocklin, H. F.; Wilson, D. M.; He, J.; Zhang, L.; Steri, V.; Wong, S. W.; Martin, T. G.; Seo, Y.; Liu, B.; Wiita, A. P.; Flavell, R. R., CD46-Targeted Theranostics for PET and 225Ac-Radiopharmaceu7cal Therapy of Mul7ple Myeloma. Clinical Cancer Research 2024, 30 (5), 1009–1021.

15. Longtine, M. S.; Shim, K.; Hoegger, M. J.; Benabdallah, N.; Abou, D. S.; Thorek, D. L. J.; Wahl, R. L., Cure of Disseminated Human Lymphoma with [<sup>225</sup>Ac]Ac-Ofatumumab in a Preclinical Model. Journal of Nuclear Medicine 2023, 64 (6), 924–931.

16. Calabretta, E.; Carlo-Stella, C., The Many Facets of CD38 in Lymphoma: From Tumor-Microenvironment Cell Interactions to Acquired Resistance to Immunotherapy. Cells 2020, 9 (4).

17. Mustafa, N.; Azaman, M. I.; Ng, G. G. K.; Chng, W. J., Molecular Determinants Underlying the Anti-Cancer Efficacy of CD38 Monoclonal Antibodies in Hematological Malignancies. Biomolecules 2022, 12 (9), 1261.

18. Herrero Alvarez, N.; Michel, A. L.; Viray, T. D.; Mayerhoefer, M. E.; Lewis, J. S., (89)Zr-DFO-Isatuximab for CD38-Targeted ImmunoPET Imaging of Multiple Myeloma and Lymphomas. ACS Omega 2023, 8 (25), 22486–22495.

19. Bauer, D.; De Gregorio, R.; Pratt, E. C.; Bell, A.; Michel, A.; Lewis, J. S., Examination of the PET in vivo generator 134Ce as a theranos7c match for 225Ac. European Journal of Nuclear Medicine and Molecular Imaging 2024.

20. Bobba, K. N.; Bidkar, A. P.; Wadhwa, A.; Meher, N.; Drona, S.; Sorlin, A. M.; Bidlingmaier, S.; Zhang, L.; Wilson, D. M.; Chan, E.; Greenland, N. Y.; Aggarwal, R.; VanBrocklin, H. F.; He, J.; Chou, J.; Seo, Y.; Liu, B.; Flavell, R. R., Development of CD46 targeted alpha theranostics in prostate cancer using <sup>134</sup>Ce/<sup>225</sup>Ac-Macropa-PEG<sub>4</sub>-YS5. TheranosBcs 2024, 14 (4), 1344–1360.

21. Rossi, M.; Botta, C.; Arbitrio, M.; Grembiale, R. D.; Tagliaferri, P.; Tassone, P., Mouse models of mul7ple myeloma: technologic plauorms and perspec7ves. Oncotarget 2018, 9 (28), 20119–20133.

22. Sgouros, G., Bone marrow dosimetry for radioimmunotherapy: theore7cal considera7ons. J Nucl Med 1993, 34 (4), 689–94.

23. Emami, B.; Lyman, J.; Brown, A.; Coia, L.; Goitein, M.; Munzenrider, J. E.; Shank, B.; Solin, L. J.; Wesson, M., Tolerance of normal 7ssue to therapeu7c irradia7on. Int J Radiat Oncol Biol Phys 1991, 21 (1), 109–22.

