## Supplementary Information for "Targeted Alpha Therapy with [^225^Ac]Ac-Macropa-Isatuximab for CD38-positive Hematological Malignancies"

### Graphical Abstract

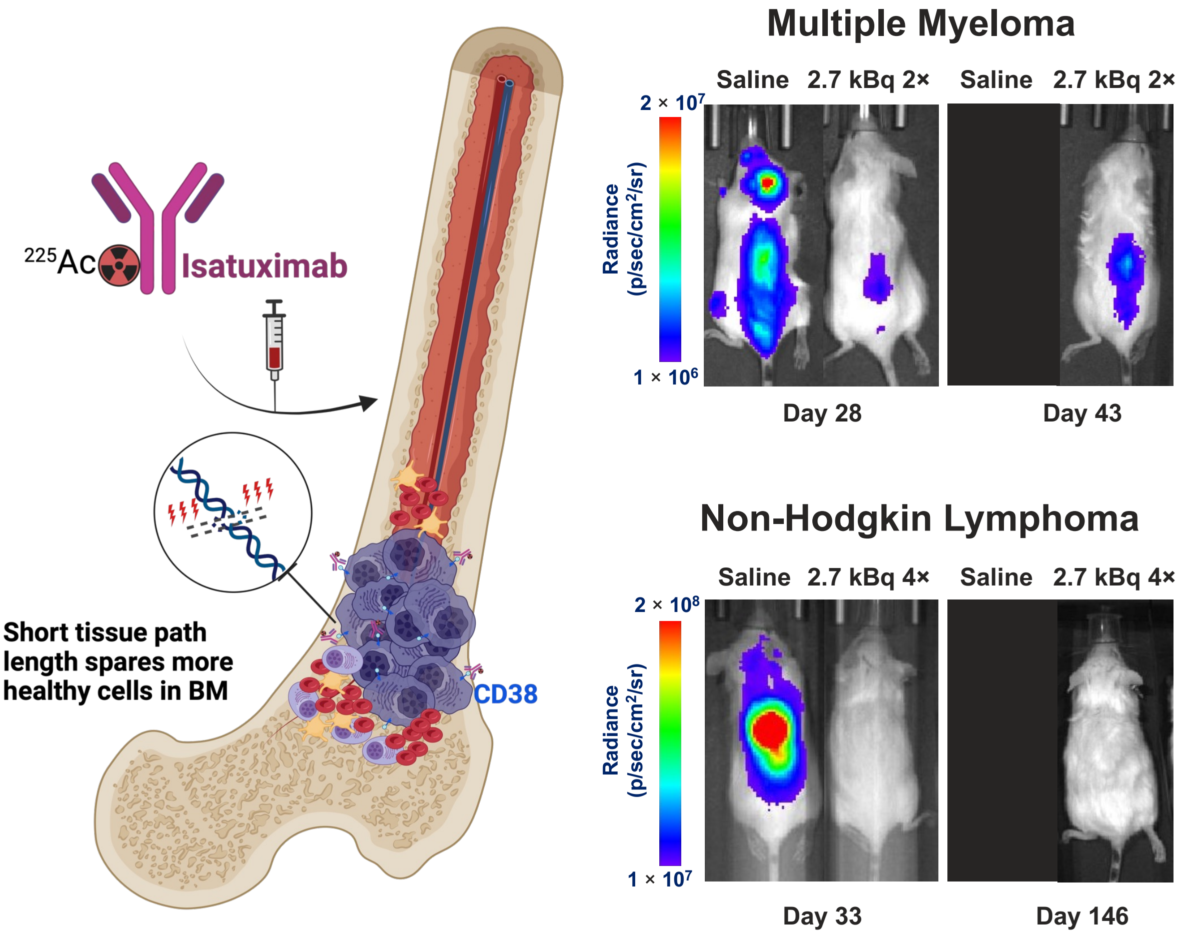

### MATERIAL AND METHODS

#### General Materials

Isatuximab (Sarclisa®; isatuximab-irfc in the USA) was purchased from the Memorial Sloan Kettering Cancer Center (MSKCC) pharmacy. Isotype human IgG was purchased from BioXCell (#BE0297). TCO-NHS ester was purchased from BroadPharm (BP-22417). $^{229}$Th-derived [$^{225}$Ac]Ac($NO_3$)$_3$ was produced and supplied by the Department of Energy isotope (DOE) program.

#### Cell Lines

MM1.S and Daudi cell lines were purchased from American Type Culture Collection (ATCC). Cultures were grown in aseptic conditions at 37°C and 5% CO2 in a humidified atmosphere. Cells were grown in RPMI-1640 medium supplemented with 20% FCS, 2 mM L-glutamine, 100 units/mL penicillin, and 100 µg/mL streptomycin. MM.1S and Daudi cells were stably transfected with firefly luciferase (Luc) through a lentivirus, allowing bioluminescence imaging (BLI). Successful transfection was confirmed by a luciferase reporter assay (Promega).

#### Immunoreactivity

Immunoreactivity was validated in a magnetic bead assay using Streptavidin magnetic beads (R&D systems) following a reported procedure.[1] Blocking was performed with a 5000-fold excess of unlabeled isatuximab. Naked beads were used as a control for unspecific binding. Measurements were run in triplicates.

#### Cell Binding Assays

Cell binding studies were performed by incubating 5 million cells with [$^{225}$Ac]Ac-Macropa-isatuximab in 1% BSA PBS at 4°C for 2 h. The blocking control contained a 100-fold excess of unlabeled isatuximab. Unbound radiotracer was removed by washing the cells twice with ice-cold PBS and centrifuging (1000 g for 5 min). Supernatant, washes, and cell pellets radioactivity was measure on a gamma counter. %Bound radioactivity was determined as the activity of the cell pellet divided by the total added activity. sed as a control for unspecific binding.

**In Vitro Stability**

In vitro stability of was [$^{225}$Ac]Ac-Macropa-isatuximab was determined by incubation in human serum at 37°C over 240 h. Demetallation was measured by iTLC and serum stability was determined as % of [$^{225}$Ac]Ac that stays chelated. Studies were performed in triplicates.

**Dosimetry**

Dosimetry calculations were performed according to the method of reference.[2] Briefly, the dosimetry estimates were computed under the following assumptions: the absorbed dose is dominated by contributions from non-penetrating radiation (e.g., alpha and beta particles) from $^{225}$Ac and its decay progeny and thus the dose contribution of photons is ignored; the progeny decay at the site of the original $^{225}$Ac decay (i.e., no redistribution of progeny); the energy deposited by alpha and beta particles is entirely deposited locally (i.e., within the organ of their origin). Using the notation of the MIRD formalism[3], the relative biological effectiveness-weighted dose coefficient (RBE-weighted dose coefficient) [Gy-equivalent per Bq $^{225}$Ac administered] for the entire decay chain is:

$$d_{RBE\text{-}weighted}\left[{}^{Gy\text{-}eq}/_{Bq}\right] = \frac{\tilde{a}}{M}\sum_i b_i \sum_j w_{R,j}\Delta_{i,j} \qquad \text{Eqn. 1}$$

where the index $i$ represents a radionuclide in the chain ($^{225}$Ac, $^{221}$Fr, $^{217}$At, $^{213}$Bi, $^{209}$Tl, $^{213}$Po, or $^{209}$Pb), the index $j$ represents the radiation type (alpha, beta, or electron), and $b_i$ is the cumulative branching fraction for chain member $i$ (1.0 [$^{225}$Ac], 1.0 [$^{221}$Fr], 1.0 [$^{217}$At], 1.0 [$^{213}$Bi], 0.022 [$^{209}$Tl], 0.978 [$^{213}$Po], 1.0 [$^{209}$Pb]), and $\Delta_{i,j}$ is the equilibrium absorbed dose constant [Gy-g/Bq-s] for the corresponding chain member and radiation type. The values of $b_i$ and $\Delta_{i,j}$ were obtained from ICRP Publication 107[4]. $w_{R,j}$ is the radiation weighting factor for *deterministic effects* for radiation type $j$, and was assumed to be 5.0 for alpha particles or 1.0 for electrons and beta particles. The quantity $\frac{\tilde{a}}{M}$ is the time-integrated activity coefficient per unit mass, where the mass is denoted $M$ [g]:

$$\frac{\tilde{a}}{M} = \int_0^{T_D} \frac{a(t)}{M} dt \qquad \text{Eqn. 2}$$

with

$$\frac{a(t)}{M} = \frac{A(t)}{M \cdot A_0} = \frac{\left[{}^{\%ID}/_{g}\right]_{bio}(t)}{100\%} \cdot e^{-\lambda_{phys} t} \qquad \text{Eqn. 3}$$

where $T_D$ is the dose integration period (taken as ∞; i.e., to complete decay), $A_0$ is the administered activity of $^{225}$Ac [Bq], $\lambda_{phys}$ is the physical half-life of $^{225}$Ac, and $\left[{}^{\%ID}/_{g}\right]_{bio}(t)$ are the organ level biodistribution values reported elsewhere in the manuscript. The integral in Eqn. 2 was evaluated by the trapezoidal method, and assumed clearance occurred by physical decay only following the last measured time point. For the purposes of dosimetry for the red bone marrow, the activity concentration of uninvolved red marrow was estimated to be 36% of that of the harvested blood samples.[5]

**Pathology**

Mice were euthanized with $CO_2$ and submitted to LCP for a comprehensive evaluation by board-certified veterinary pathologists. Organs were fixed in 10% neutral buffered formalin, followed by decalcification of the bones and other hard tissues in a formic acid solution (Surgipath Decalcifier I, Leica Biosystems). Tissues were processed in ethanol and xylene and embedded in paraffin in a Leica ASP6025 tissue processor. Paraffin blocks were sectioned at 5 microns and stained with hematoxylin and eosin (H&E).

**Immunohistochemistry**

The immunohistochemistry detection of CD38 was performed at Molecular Cytology Core Facility of Memorial Sloan Kettering Cancer Center, using Discovery XT processor

(Ventana Medical Systems, Roche - AZ). Femurs were decalcified prior to sectioning. After 32 min of heat and CC1 (Cell Conditioning 1, Ventana catalog#: 950-500) retrieval, the tissue sections were blocked first for 30 min in Background Blocking reagent (Innovex, catalog#: NB306). A rabbit monoclonal anti-CD38 antibody (Ventana Ref#: 790-7016) was used in prediluted ready-to-use concentration. The incubation with the primary antibody was done for 4 h, followed by 60 min incubation with biotinylated goat anti-rabbit IgG (Vector labs, catalog#: PK6101) in 5.75 µg/mL, followed by application of Blocker D, Streptavidin- HRP and DAB detection kit (Ventana Medical Systems), according to the manufacturer instructions.

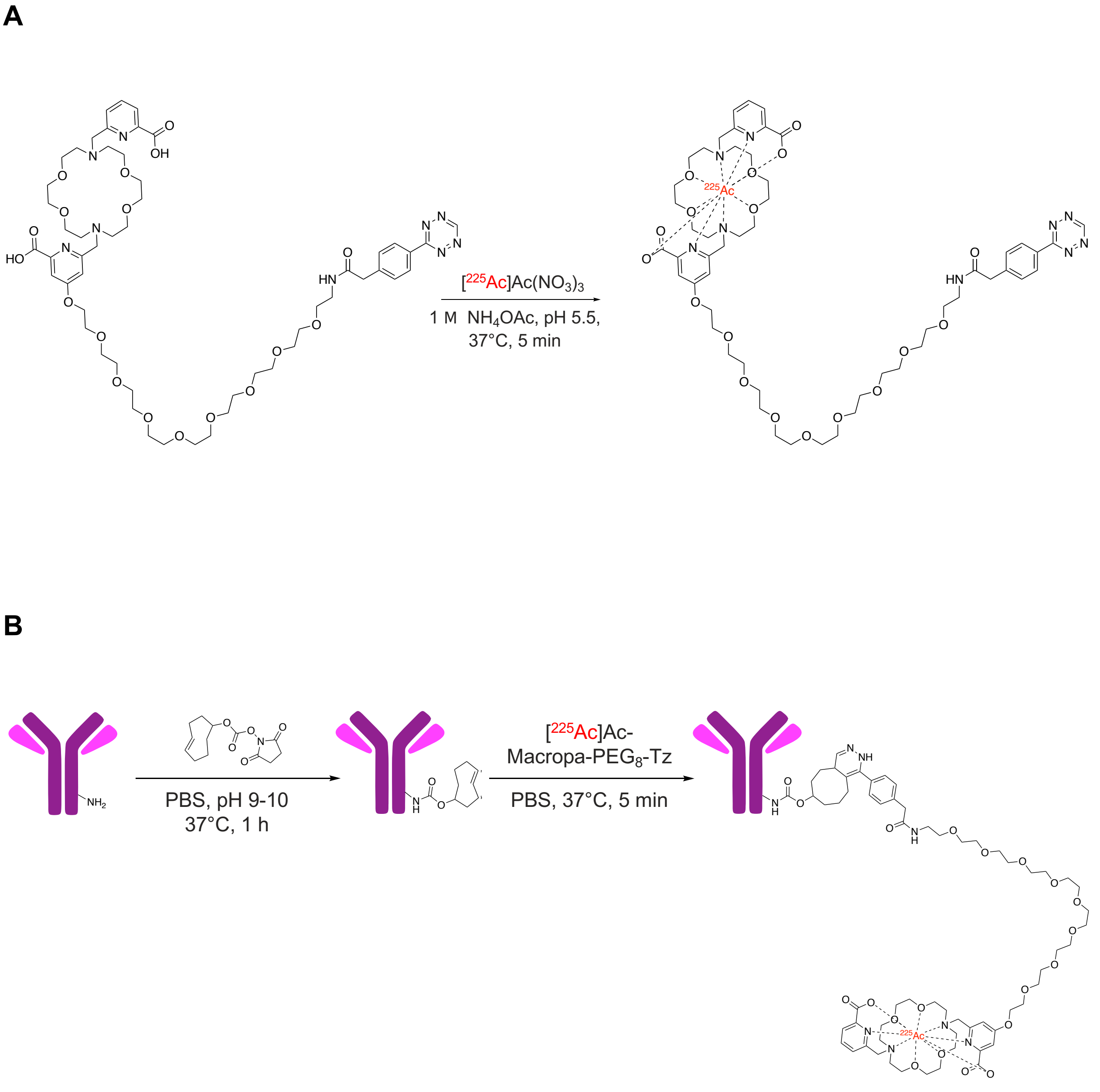


**Supplemental Figure 1. Synthesis overview of [$^{225}$Ac]Ac-Macropa-PEG$_8$-isatuximab.** Radiolabeling of Macropa-PEG$_8$-Tz with $^{225}$Ac (A). Bioconjugation of isatuximab to the TCO moiety, followed by Inverse Electron-Demand Diels Alder click reaction with [$^{225}$Ac]Ac-Macropa-PEG$_8$-Tz (B).

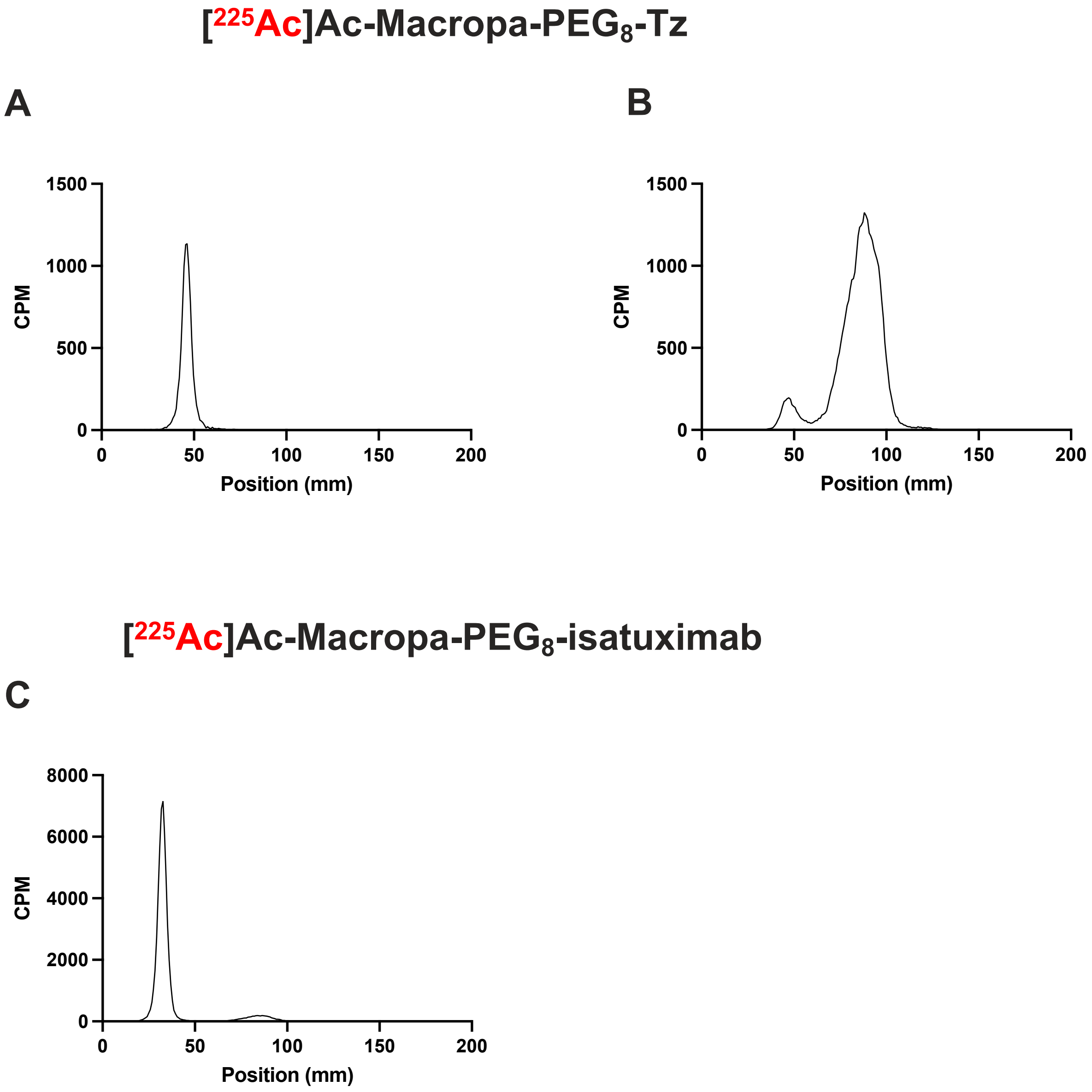


**Supplemental Figure 2.** Radio-instant thin layer chromatography of [$^{225}$Ac]Ac-Macropa-PEG$_8$-Tz using 50 mM ethylenediaminetetraacetic acid (A) and 0.25 M $NH_4OAc$/methanol (40/60) as the mobile phase (B) showing high radiochemical conversion > 99%. Radio-instant thin layer chromatography of [$^{225}$Ac]Ac-Macropa-PEG$_8$-isatuximab using 0.25 M $NH_4OAc$/methanol (40/60) as the mobile phase (C) proving completion of the click reaction > 95%.

**A**

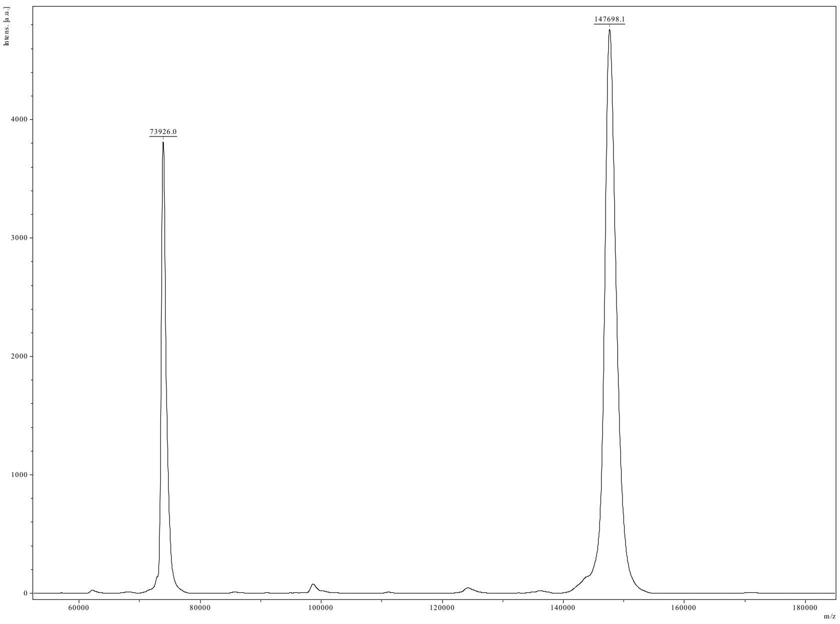


**B**

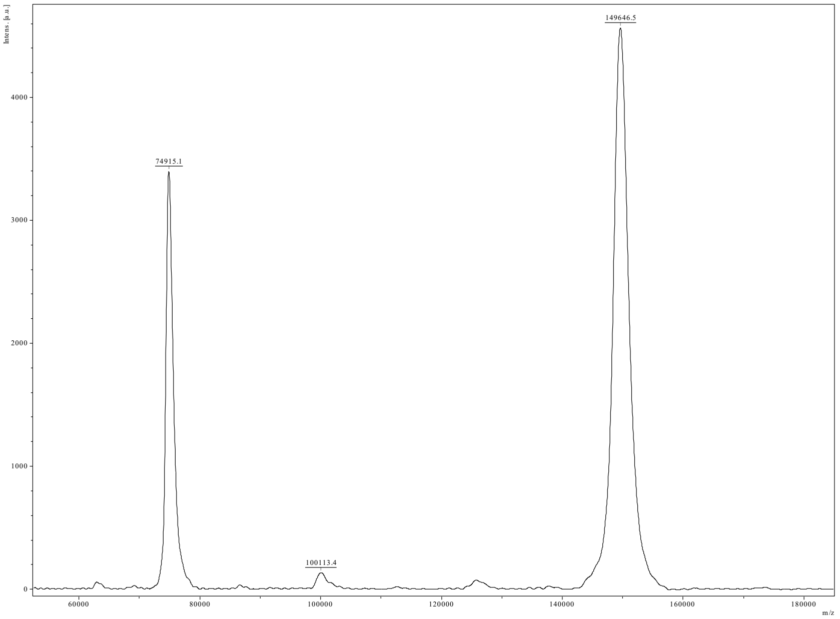


**Supplemental Figure 3.** MALDI-TOF spectra of native and modified isatuximab. Native isatuximab shows a peak at MW 147698 (A) and isatuximab-TCO at MW 149646 (B), resulting in an average of 7 TCO moieties per molecule of antibody.

### In Vitro Evaluation

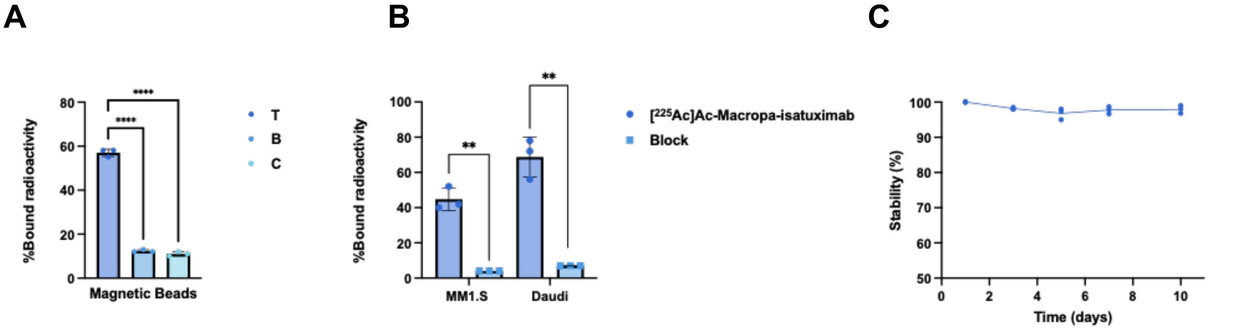


**Supplemental Figure 4. *In vitro* characterization of [$^{225}$Ac]Ac-Macropa-isatuximab.** CD38 bead-binding assay shows high and specific binding of [$^{225}$Ac]Ac-Macropa-isatuximab to the receptor. Binding is blockable with a 1000-fold excess of unlabeled isatuximab (A). Cell binding assays show high and specific binding of [$^{225}$Ac]Ac-Macropa-isatuximab to MM1.S and Daudi cell lines. Binding is blockable with a 100-fold excess of unlabeled isatuximab (B). [$^{225}$Ac]Ac-Macropa-isatuximab shows excellent stability >98% in human serum over 10 days at 37°C.

A

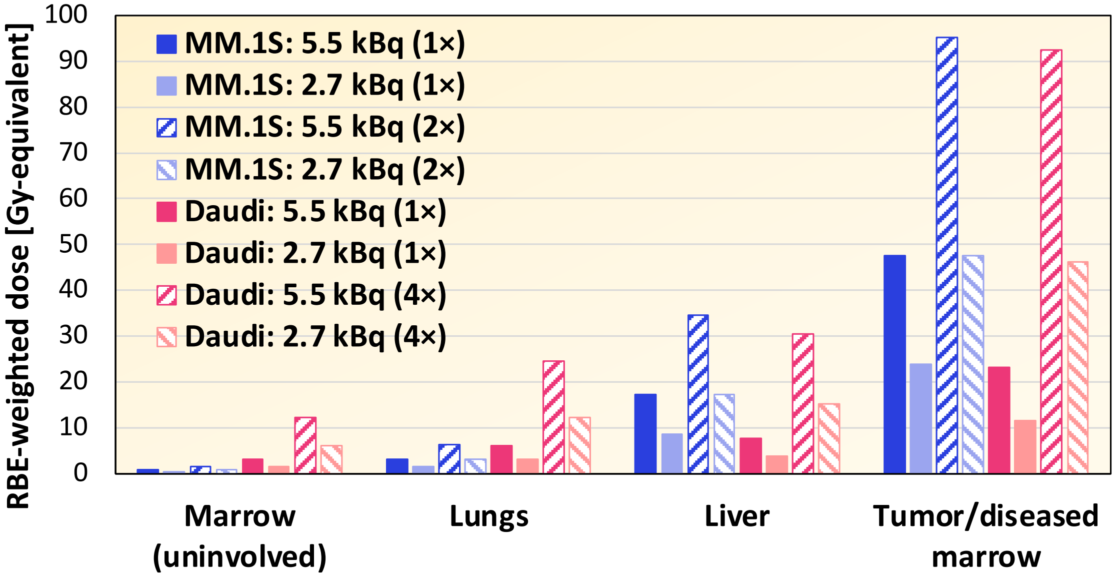

RBE-weighted dose [Gy-equivalent]
100
90
80
70
60
50
40
30
20
10
0
MM.1S: 5.5 kBq (1×)
MM.1S: 2.7 kBq (1×)
MM.1S: 5.5 kBq (2×)
MM.1S: 2.7 kBq (2×)
Daudi: 5.5 kBq (1×)
Daudi: 2.7 kBq (1×)
Daudi: 5.5 kBq (4×)
Daudi: 2.7 kBq (4×)
Marrow (uninvolved)
Lungs
Liver
Tumor/diseased marrow

**B**

**MM.1s**

| Organ | RBE-weighted dose coefficient (Gy-equivalent/uCi [$^{225}$Ac]Ac-mcp-Isatuximab) | TI |
|---|---|---|
| Marrow (uninvolved) | 5.14 | 62.6 |
| Heart | 7.02 | 45.9 |
| Lungs | 21.6 | 14.9 |
| Liver | 117 | 2.7 |
| Pancreas | 5.72 | 56.3 |
| Stomach | 3.60 | 89.5 |
| Small Intestine | 4.62 | 69.7 |
| Colon | 3.66 | 88.0 |
| Kidneys | 37.4 | 8.6 |
| Muscle | 5.13 | 62.7 |
| Bone | 239 | 1.3 |
| Tumor/diseased marrow | 322 | -- |

**Daudi**

| Organ | RBE-weighted dose coefficient (Gy-equivalent/uCi [$^{225}$Ac]Ac-mcp-Isatuximab) | TI |
|---|---|---|
| Marrow (uninvolved) | 20.8 | 7.5 |
| Lungs | 41.3 | 3.8 |
| Liver | 51.5 | 3.0 |
| Tumor/diseased marrow | 156 | -- |

**Supplemental Figure 5.** Calculated organ-level absorbed doses in MM1.S and Daudi xenografts using [$^{89}$Zr]Zr-DFO-Isatuximab biodistribution data (**A**). Tabulated absorbed dose values and therapeutic index for the different cohorts (**B**).

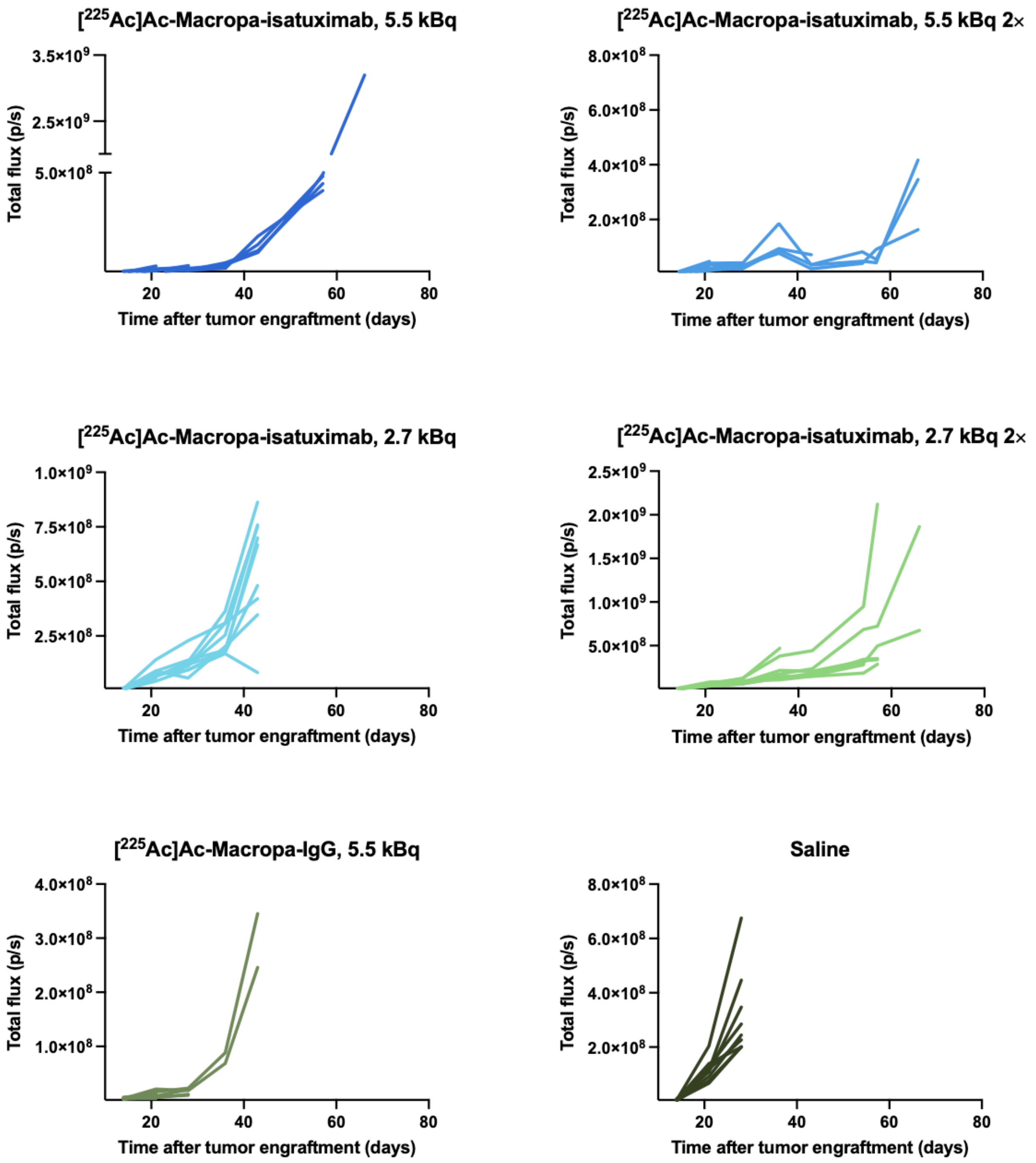


**Supplemental Figure 6.** Individual mouse tumor burden quantification by BLI for NSG mice with MM1.S disseminated disease for all the cohorts of the therapy study.

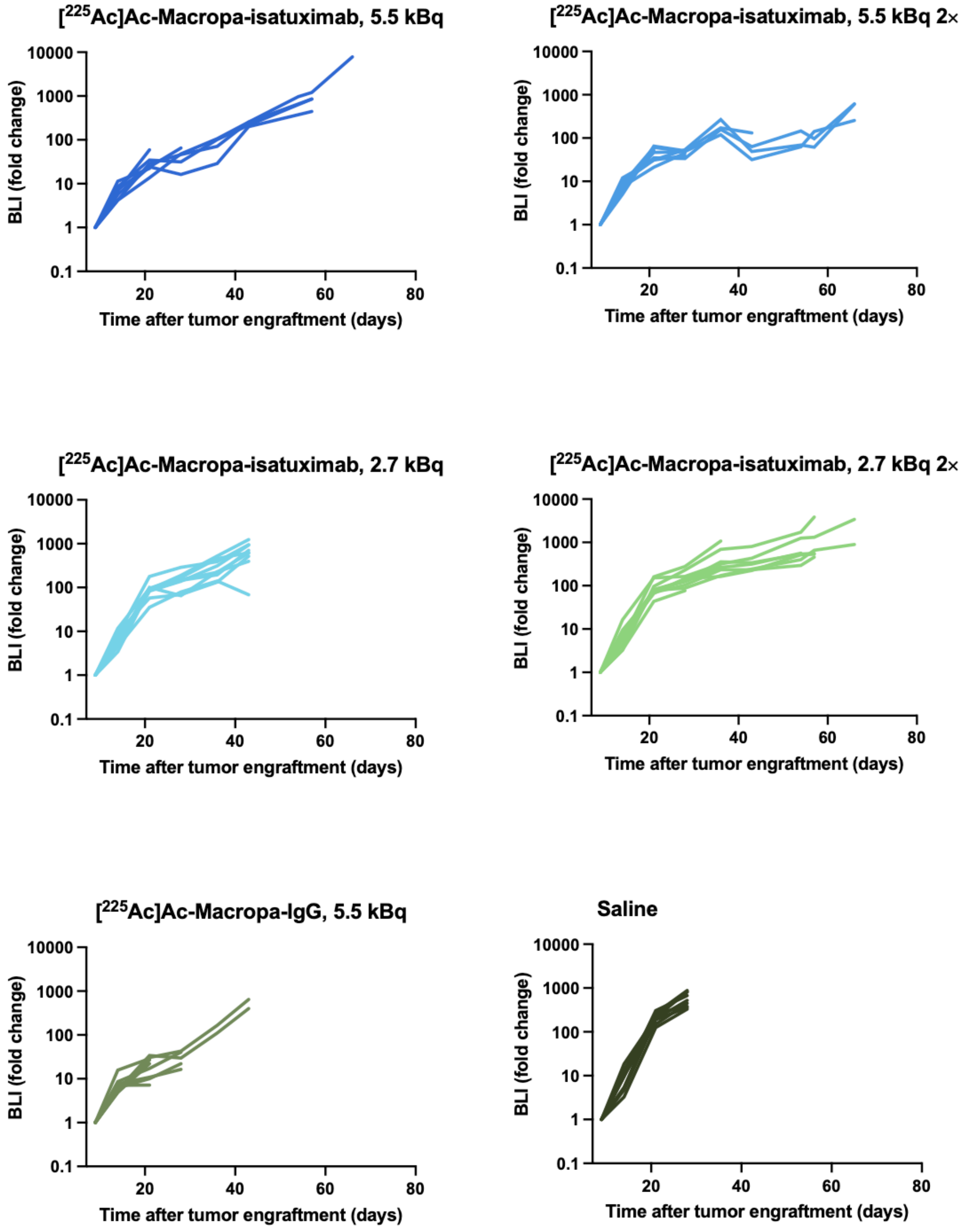


**Supplemental Figure 7.** Individual mouse tumor burden fold change measured by BLI for NSG mice with MM1.S disseminated disease for all the cohorts of the therapy study.

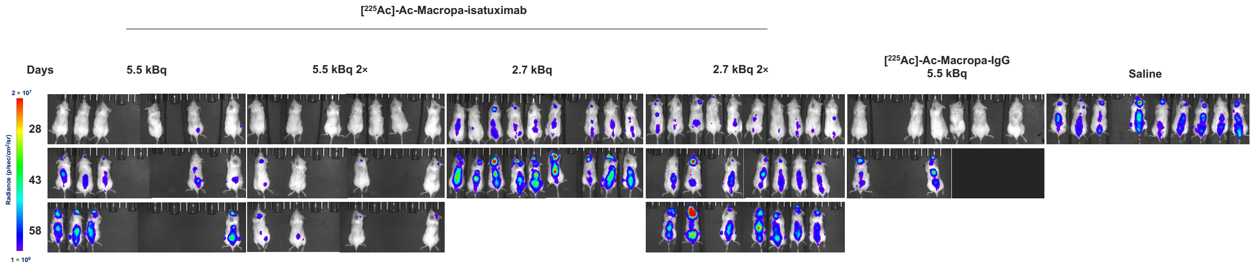


**Supplemental Figure 8.** BLI of NSG mice with MM1.S disseminated disease for all the cohorts of the therapy study at the most representative days.

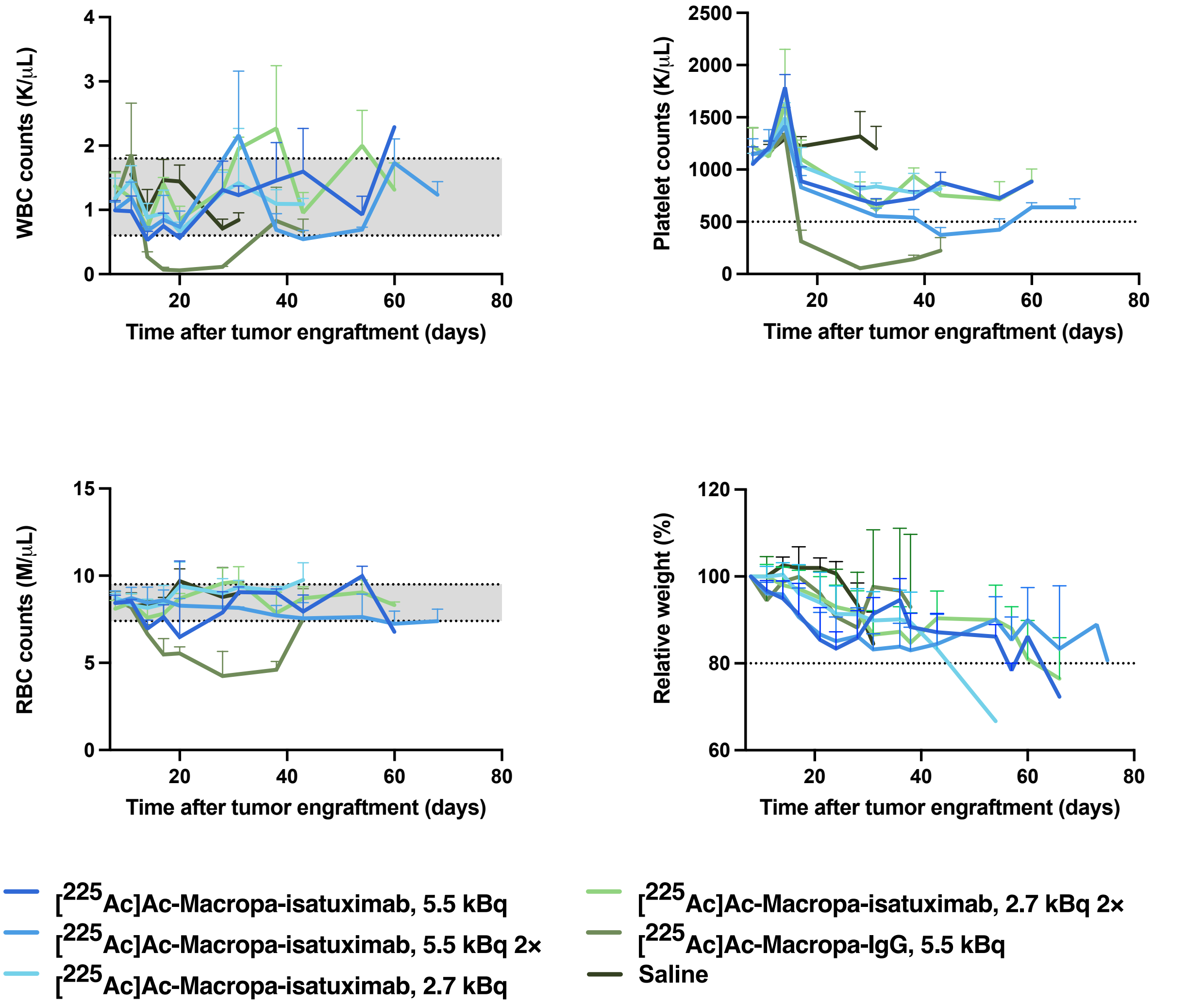


**Supplemental Figure 9.** Hematologic toxicity is transitory upon treatment with [$^{225}$Ac]Ac-Macropa-isatuximab and normal values are recovered after a few weeks (A–C). White blood cell counts (A), platelet counts (B), and red cell counts (C) in MM1.S disseminated NSG mice. Shading indicates mean ± SD of pretherapy values (week 0). (D) Relative weight percentages for MM1.S disseminated mice in each treatment cohort. WBC = white blood cell; RBC = red blood cell.

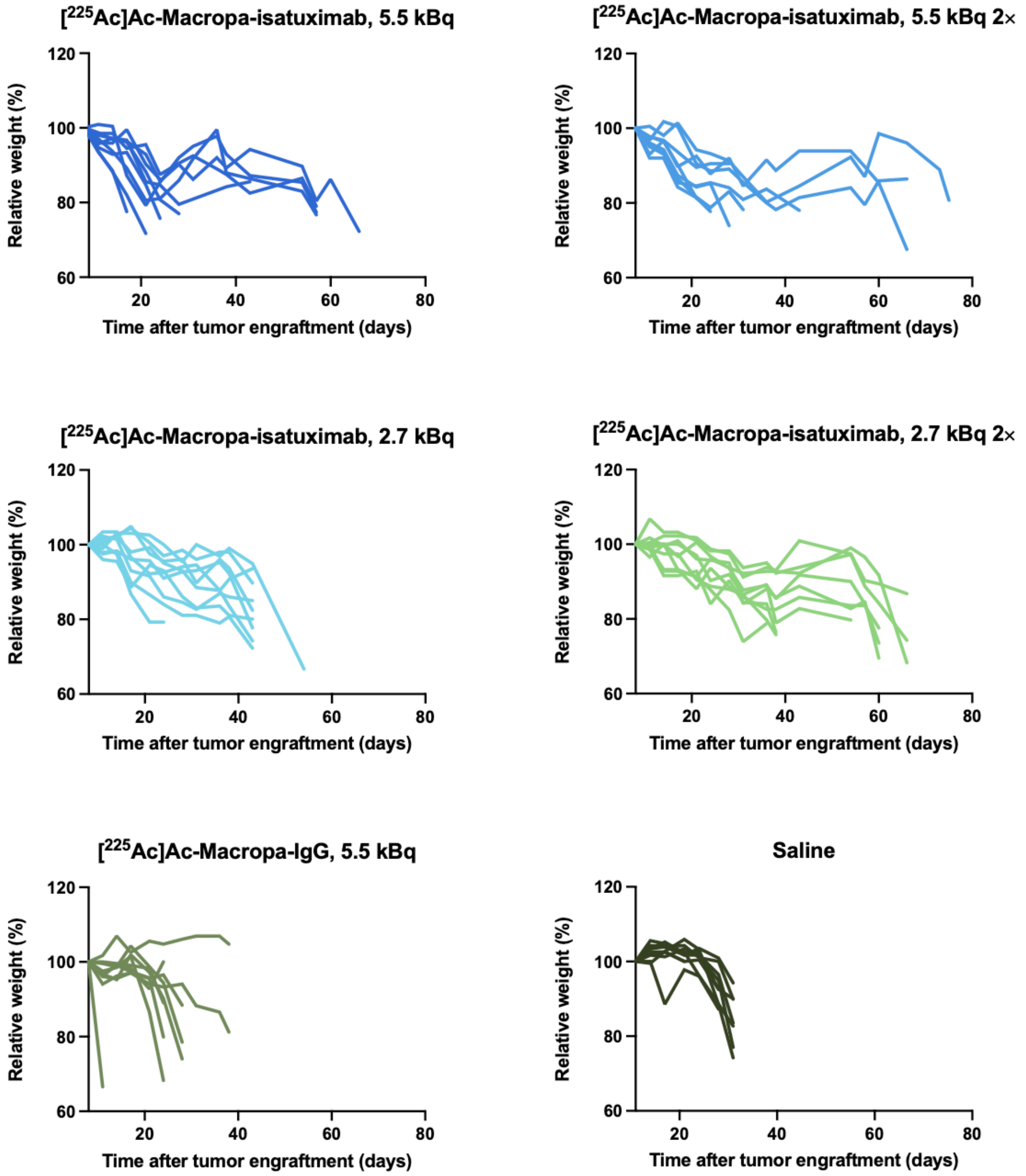


**Supplemental Figure 10.** Individual mouse relative weight percentages for NSG mice with MM1.S disseminated disease for all the cohorts of the therapy study.

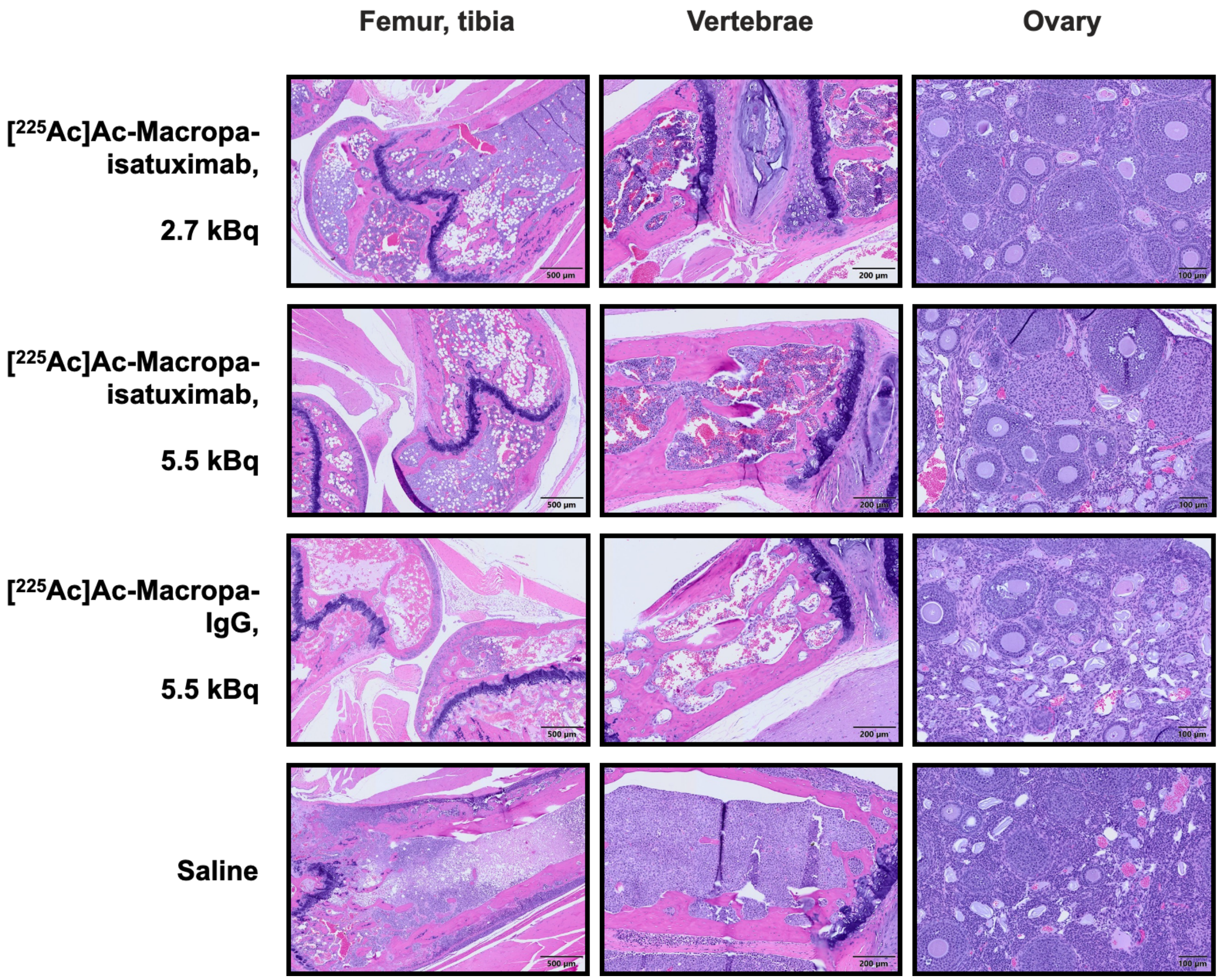


**Supplemental Figure 11.** Representative histology of hematoxylin- and eosin-stained femur/tibia (left), vertebrae (middle), and ovarian sections of MM1.S disseminated NSG mice. Therapeutic effect is dose-dependent as seen by a more profound tumor growth inhibition in mice treated with 5.5 kBq [$^{225}$Ac]Ac-Macropa-isatuximab compared to mice treated with 2.7 kBq [$^{225}$Ac]Ac-Macropa-isatuximab. Mice treated with the lower dose (first row) presented moderated to marked osteolysis and marked infiltration of neoplastic cells into femur, tibia, and vertebrae. Mice treated with the higher dose (second row) presented mild osteolysis and mild to moderate infiltration of neoplastic cells into femur and vertebrae. Ovaries showed diffuse mild atrophy in both groups. Untargeted .5 kBq [$^{225}$Ac]Ac-Macropa-IgG presented moderate osteolysis, marked hypoplasia, and hemorrhage as well as moderate infiltration of neoplastic cells into vertebrae. Toxicity to the ovaries was also higher in this cohort, showing moderate atrophy. Finally, the untreated mice (last row) presented marked osteolysis and infiltration of neoplastic cells into femur, tibia, and vertebrae with widespread necrosis and hemorrhage. Infiltration of tumor cells reached external musculature and the meninges resulting in spinal cord compression, showing the aggressiveness of the disease when untreated. Histopathology of the ovaries was normal.

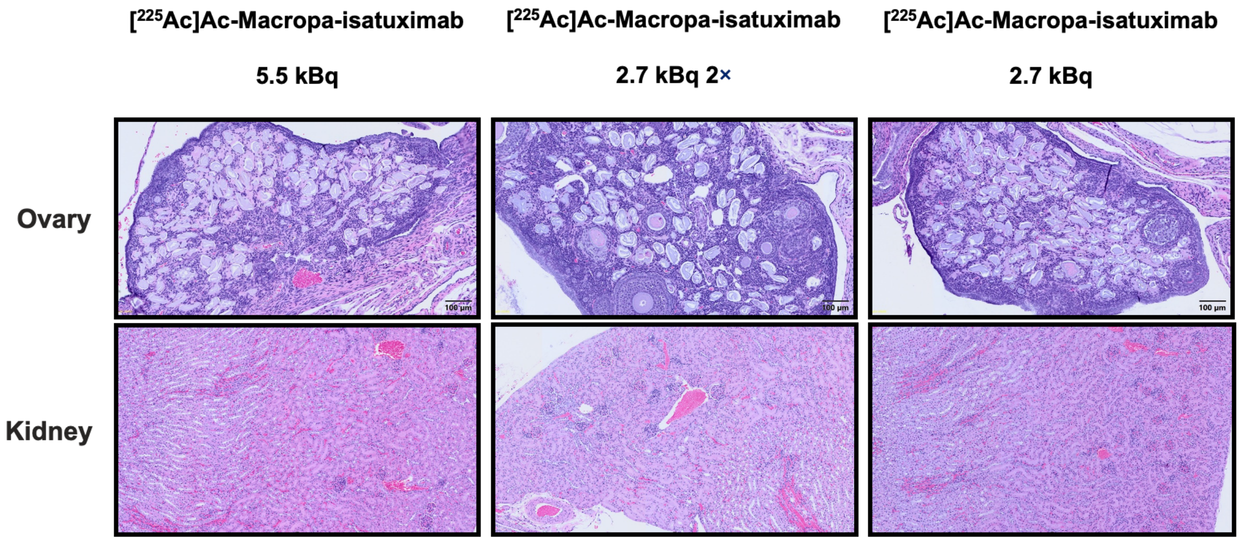


**Supplemental Figure 12.** Representative histology of hematoxylin- and eosin-stained ovarian (top) and kidney (bottom) sections of MM1.S disseminated NSG mice. Histopathology of the ovaries revealed diffuse atrophy with loss of follicles. Degree of atrophy was marked for mice treated with 5.5 kBq [$^{225}$Ac]Ac-Macropa-isatuximab (left) and 2.7 kBq [$^{225}$Ac]Ac-Macropa-isatuximab 2× (middle), and moderate for mice treated with 2.7 kBq [$^{225}$Ac]Ac-Macropa-isatuximab (right). Histopathology of kidneys revealed a minimal multifocal tubular cast formation for mice treated with 5.5 kBq [$^{225}$Ac]Ac-Macropa-isatuximab (left) and minimal focal tubular attenuation for mice treated with 2.7 kBq [$^{225}$Ac]Ac-Macropa-isatuximab 2× (middle). Kidneys of mice treated with 2.7 kBq [$^{225}$Ac]Ac-Macropa-isatuximab (right) remained histologically normal, showing that this organ is minimally affected by the therapy.

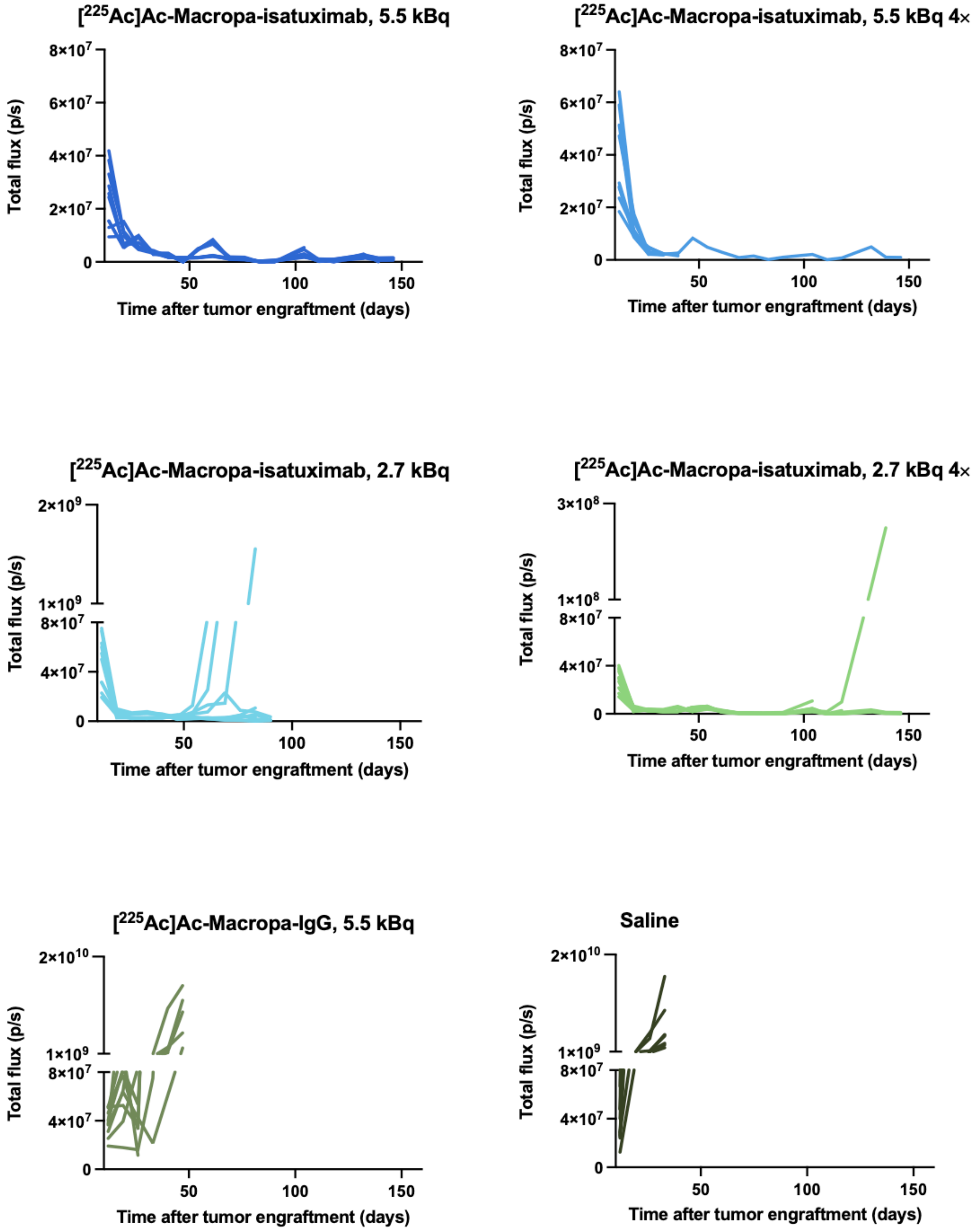


**Supplemental Figure 13.** Individual mouse tumor burden quantification by BLI for NSG mice with Daudi disseminated disease for all the cohorts of the therapy study.

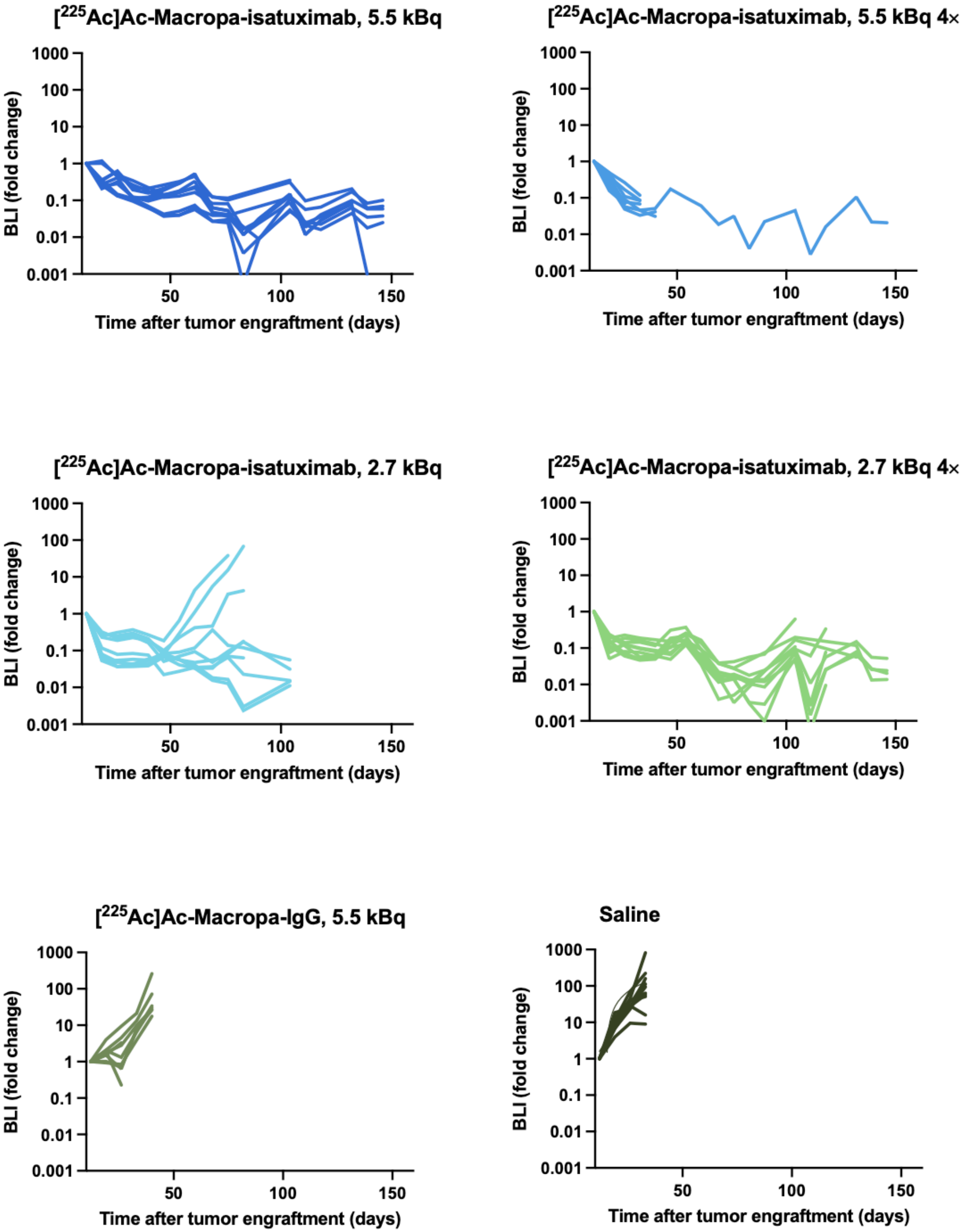


**Supplemental Figure 14.** Individual mouse tumor burden fold change measured by BLI for NSG mice with Daudi disseminated disease for all the cohorts of the therapy study.

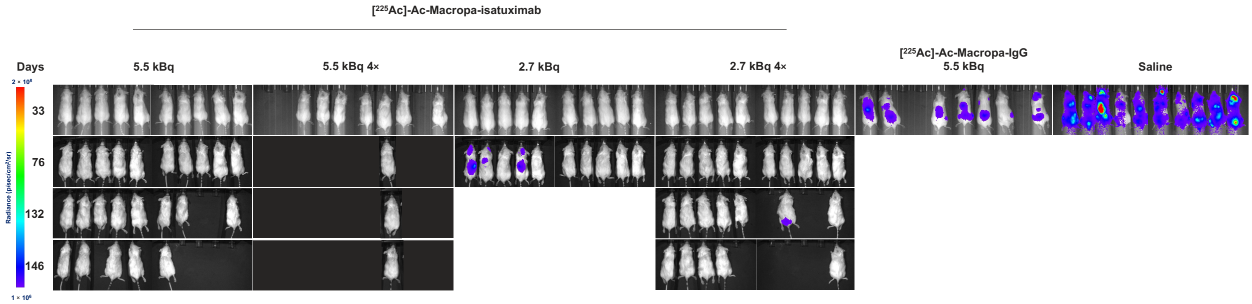


**Supplemental Figure 15.** BLI of NSG mice with Daudi disseminated disease for all the cohorts of the therapy study at the most representative days.

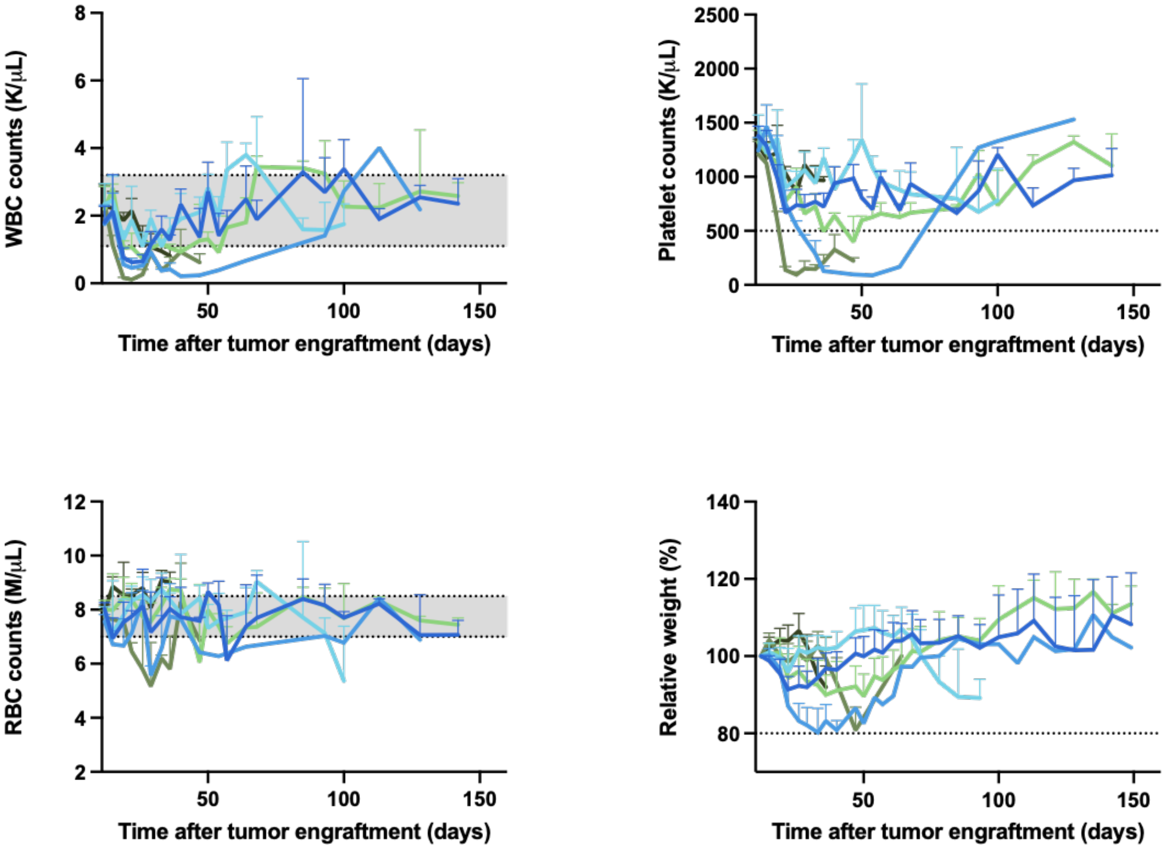


**Supplemental Figure 16.** Hematologic toxicity is transitory upon treatment with [$^{225}$Ac]Ac-Macropa-isatuximab and normal values are recovered after a few weeks (A-C). White blood cell counts (A), platelet counts (B), and red cell counts (C) in Daudi disseminated NSG mice. Shading indicates mean ± SD of pretherapy values (week 0). (D) Relative weight percentages for Daudi disseminated mice in each treatment cohort. WBC = white blood cell; RBC = red blood cell.

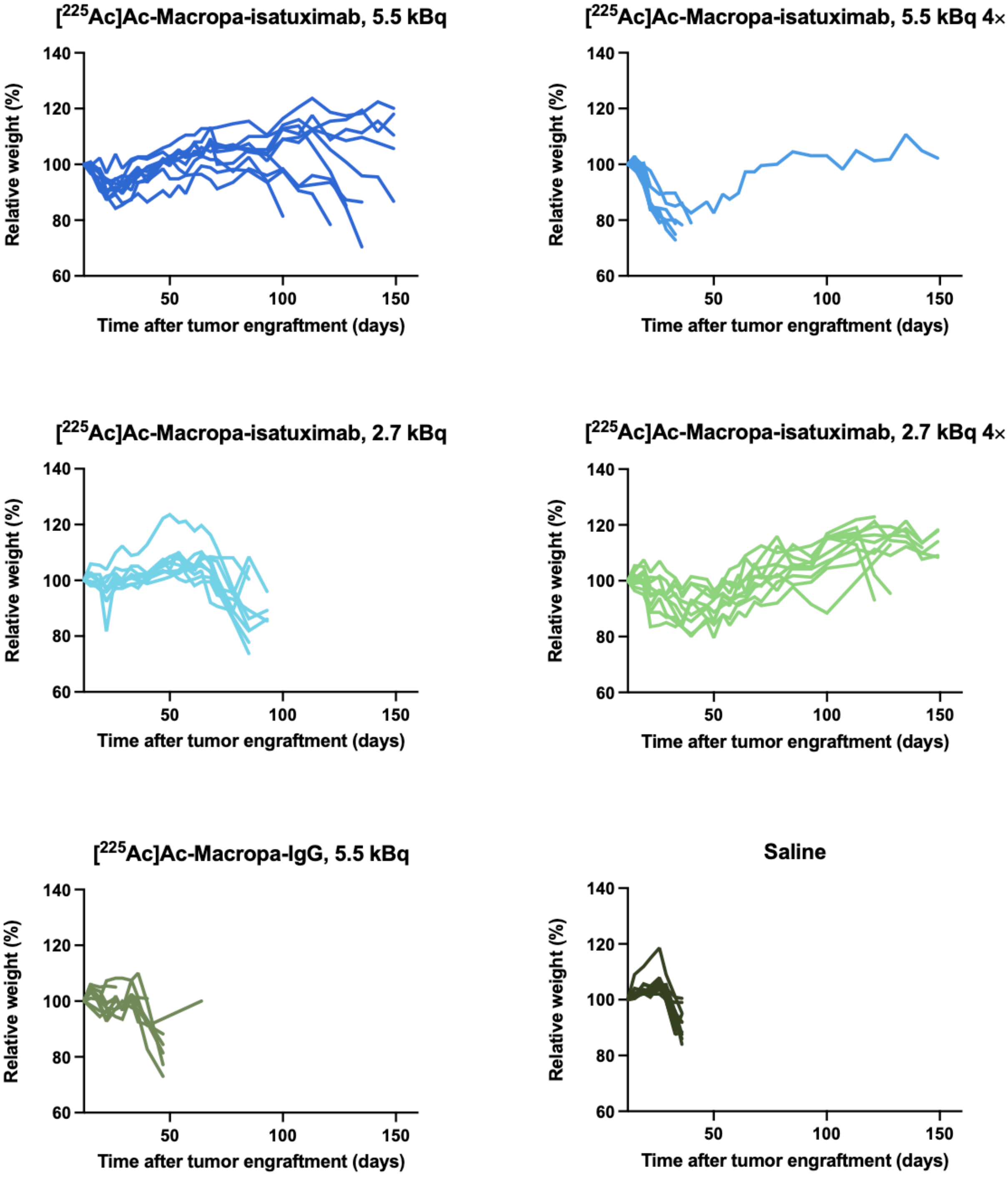


**Supplemental Figure 17.** Individual mouse relative weight percentages for NSG mice with Daudi disseminated disease for all the cohorts of the therapy study.

**Comparative Pathology - Daudi**

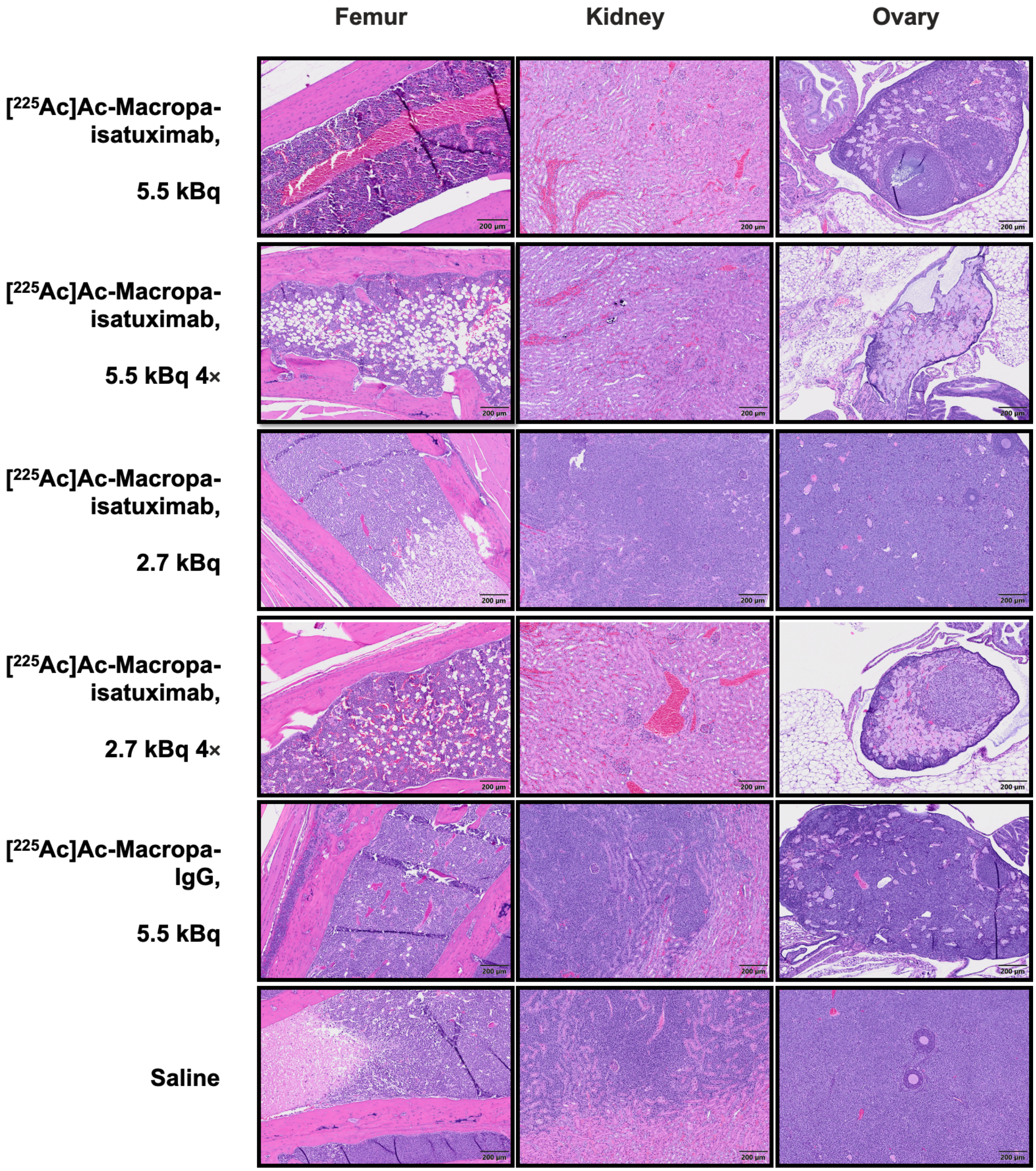


**Supplemental Figure 18.** Representative histology of hematoxylin- and eosin-stained femur (left), kidneys (middle), and ovarian sections of Daudi disseminated NSG mice. A number of mice receiving [$^{225}$Ac]Ac-Macropa-isatuximab treatments with 5.5 kBq, 5.5 kBq

4×, and 2.7 kBq 4× doses reached the end of the study without tumor relapse. Optimal treatment cohorts, 5.5 kBq and 2.7 kBq 4×, presented a completely healthy bone marrow composition, while the mouse receiving the highest radiation, 5.5 kBq 4×, showed minimal to mild hypoplasia. These animals did not present any abnormality in the kidneys, demonstrating that this treatment is not associated with adverse renal events. On the other hand, they showed mild to marked ovarian atrophy which is considered a direct result of the treatment. Mice that received the lowest dose of radiation, 2.7 kBq, presented multifocal round cell infiltration in the femur, kidneys, and ovaries. Mice from the untargeted [$^{225}$Ac]Ac-Macropa-IgG cohort presented multifocal neoplastic red infiltration in the bone marrow as well as moderate hypoplasia, and severe tumor invasion in the kidneys and ovaries. Saline control mice presented neoplastic red infiltration with frequent tumor necrosis in the bone marrow, and severe tumor invasion in the kidneys and ovaries, demonstrating the aggressivity of this lymphoma model.

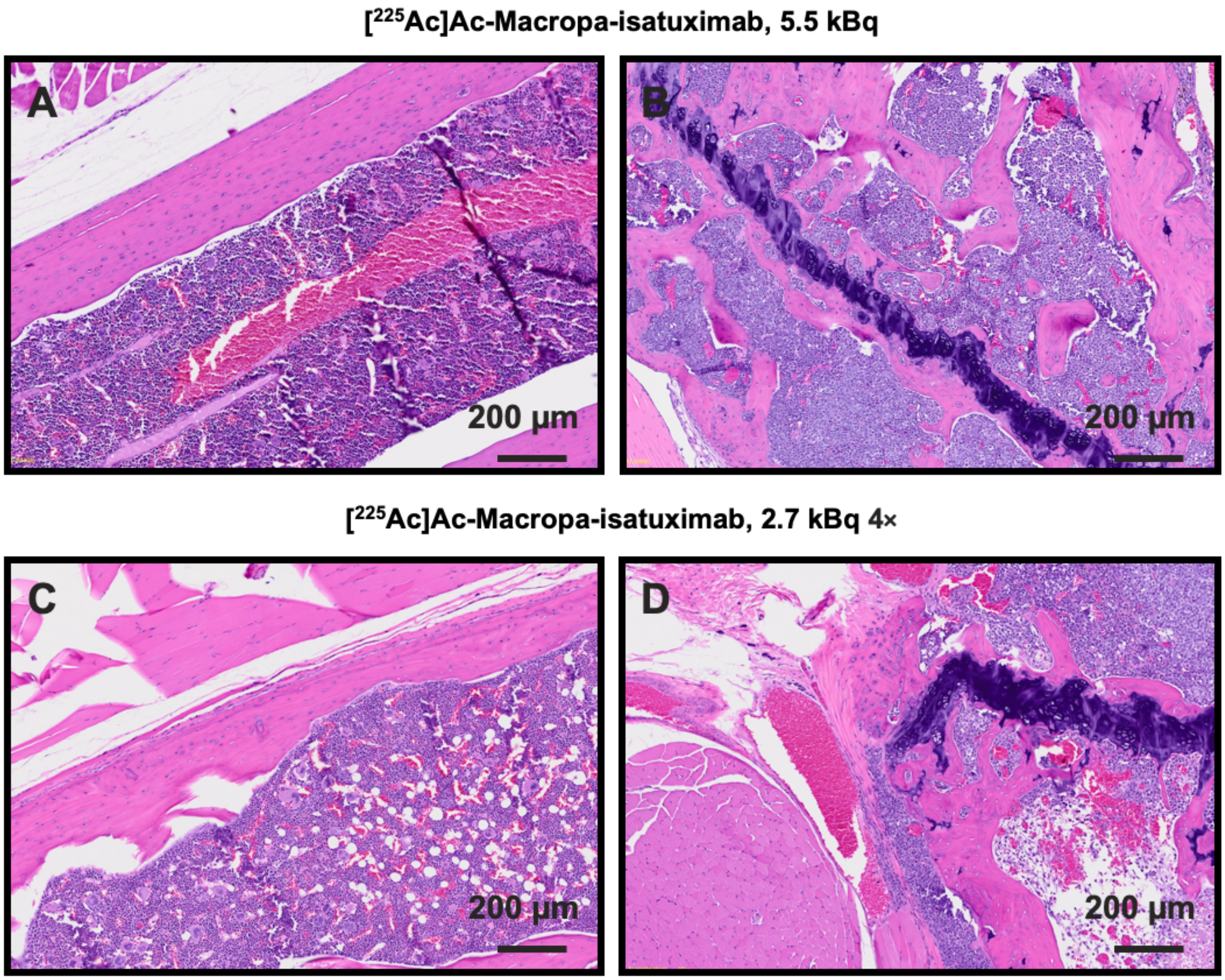


**Supplemental Figure 19.** Representative histology of hematoxylin- and eosin-stained femur sections of Daudi disseminated NSG mice from the most successful [$^{225}$Ac]Ac-Macropa-isatuximab treatment cohorts: 5.5 kBq (A,B), and 2.7 kBq 4× (C,D). The 4 mice were euthanized at the end of the study (150 days) without signs of disease by BLI. However, while the histopathology analysis demonstrated that A and C were complete responders, tumor burden was detected in B and D, suggesting a tumor relapse by Daudi cells no longer expressing luciferase. Animal B presented neoplastic round cell infiltration, multifocal hypoplasia, and moderate necrosis and hemorrhage. Animal D presented neoplastic round cell infiltration with frequent tumor necrosis.

### References


1. Sharma, S. K.; Lyashchenko, S. K.; Park, H. A.; Pillarsetty, N.; Roux, Y.; Wu, J.; Poty, S.; Tully, K. M.; Poirier, J. T.; Lewis, J. S., A rapid bead-based radioligand binding assay for the determination of target-binding fraction and quality control of radiopharmaceuticals. *Nucl Med Biol* **2019,** *71*, 32-38.
2. Poty, S.; Carter, L. M.; Mandleywala, K.; Membreno, R.; Abdel-Atti, D.; Ragupathi, A.; Scholz, W. W.; Zeglis, B. M.; Lewis, J. S., Leveraging Bioorthogonal Click Chemistry to Improve (225)Ac-Radioimmunotherapy of Pancreatic Ductal Adenocarcinoma. *Clin Cancer Res* **2019,** *25* (2), 868-880.
3. Bolch, W. E.; Eckerman, K. F.; Sgouros, G.; Thomas, S. R., MIRD pamphlet No. 21: a generalized schema for radiopharmaceutical dosimetry--standardization of nomenclature. *J Nucl Med* **2009,** *50* (3), 477-84.
4. Eckerman, K.; Endo, A., ICRP Publication 107. Nuclear decay data for dosimetric calculations. *Ann ICRP* **2008,** *38* (3), 7-96.
5. Sgouros, G., Bone marrow dosimetry for radioimmunotherapy: theoretical considerations. *J Nucl Med* **1993,** *34* (4), 689-94.
